# Pepper RNA variants reveal decoupling of HBC530 binding thermodynamics and fluorescence activation

**DOI:** 10.1101/2025.06.24.660466

**Authors:** Emmanuel K. Aidoo, Vishakha Jayasekera, Nkosinathi T. Dlamini, Patrick H. Dussault, Catherine D. Eichhorn

## Abstract

Fluorogenic RNA aptamers have emerged as powerful tools for live-cell imaging and synthetic biology applications due to their ability to activate fluorescence upon ligand binding. However, the sequence-structure-function relationships governing ligand recognition and fluorescence activation remain poorly understood, limiting rational aptamer design. The Pepper aptamer binds HBC530 with nanomolar affinity in a magnesium-dependent manner, producing bright fluorescence suitable for cellular applications. Here, we generated a library of 53 Pepper variants containing substitutions, insertions, and/or deletions to quantitatively evaluate the contributions of individual nucleotides to HBC530 binding affinity, magnesium binding affinity, and fluorescence intensity. Our results reveal that the correlation between HBC530 binding affinity and fluorescence intensity is only modest. We identify several variants with binding affinities similar to wild-type Pepper but dramatically reduced fluorescence, indicating that ligand recognition and fluorescence intensity are decoupled. Further, we find that distal structural elements significantly influence both binding thermodynamics and fluorescence intensity. Confirming the critical role of divalent cations in Pepper-HBC530 recognition, we observe a strong correlation between HBC530 and magnesium binding thermodynamics. These findings provide quantitative insights into the molecular mechanisms underlying fluorescence activation, toward developing a framework for the rational design of the next generation of fluorogenic RNA aptamers with enhanced performance.

**KEY POINTS:**

- The HBC530 binding affinities of 53 Pepper RNA variants are only moderately correlated with fluorescence intensity
- Structural elements distal to the HBC530 binding pocket influence both binding affinity and fluorescence
- The binding affinities of Pepper RNA to HBC530 and Mg^2+^ are highly correlated, and Mg^2+^ stoichiometry is correlated with binding thermodynamics and fluorescence intensity

## INTRODUCTION

Over the past two decades, multiple classes of synthetic fluorogenic RNA aptamers have been developed that bind a diverse palette of dyes with high affinity and selectivity (1–3). These dyes have minimal fluorescence in the unbound state but become highly fluorescent when bound to their cognate RNA aptamer, providing effective “light-up” reporter systems for synthetic biology and live cell imaging applications (1,4). Current methods to generate aptamers use selection-based techniques, often including magnesium (Mg^2+^) (5), to identify sequences that bind a target ligand (6–9). In addition to binding selectivity, incorporating functional properties such as fluorescence enhancement into screening criteria have resulted in improved aptamer-dye pairs with enhanced photophysical properties (10,11). While structural insights into fluorogenic aptamers have advanced a mechanistic understanding of aptamer-dye recognition, the ability to rationally design or improve aptamer-dye photophysical properties remains limited by an incomplete understanding of how RNA sequence and structure influence Mg^2+^ coordination, dye recognition, and fluorescence activation.

The fluorogenic Pepper RNA aptamer forms a monomeric complex with HBC530 ((4-((2-hydroxyethyl)(methyl)amino)-benzylidene)-cyanophenyl-acetonitrile), along with a suite of HBC derivatives, with nanomolar affinity (12). Recent X-ray crystal structures revealed the structural basis for Pepper-HBC recognition, where two internal loops (J1/2 and J2/3) form tertiary interactions to create the HBC binding pocket (**Fig. 1**) (13,14). Pepper-HBC complex formation and subsequent fluorescence activation requires Mg^2+^ (12–14). Mg^2+^ is a critical divalent cation that promotes RNA aptamer-dye complex formation by coordinating with residues through diffuse electrostatic and/or site-specific interactions to stabilize tertiary contacts and noncanonical base-pairs (15). HBC530 is soluble, photostable, non-cytotoxic, and cell permeable (12), making Pepper-HBC attractive for biological applications. Despite increasing adoption of Pepper in biosensing and imaging applications (16–20) quantitative understanding of the sequence determinants governing HBC530 recognition, magnesium coordination, and fluorescence activation remains incomplete.

**Figure 1.**
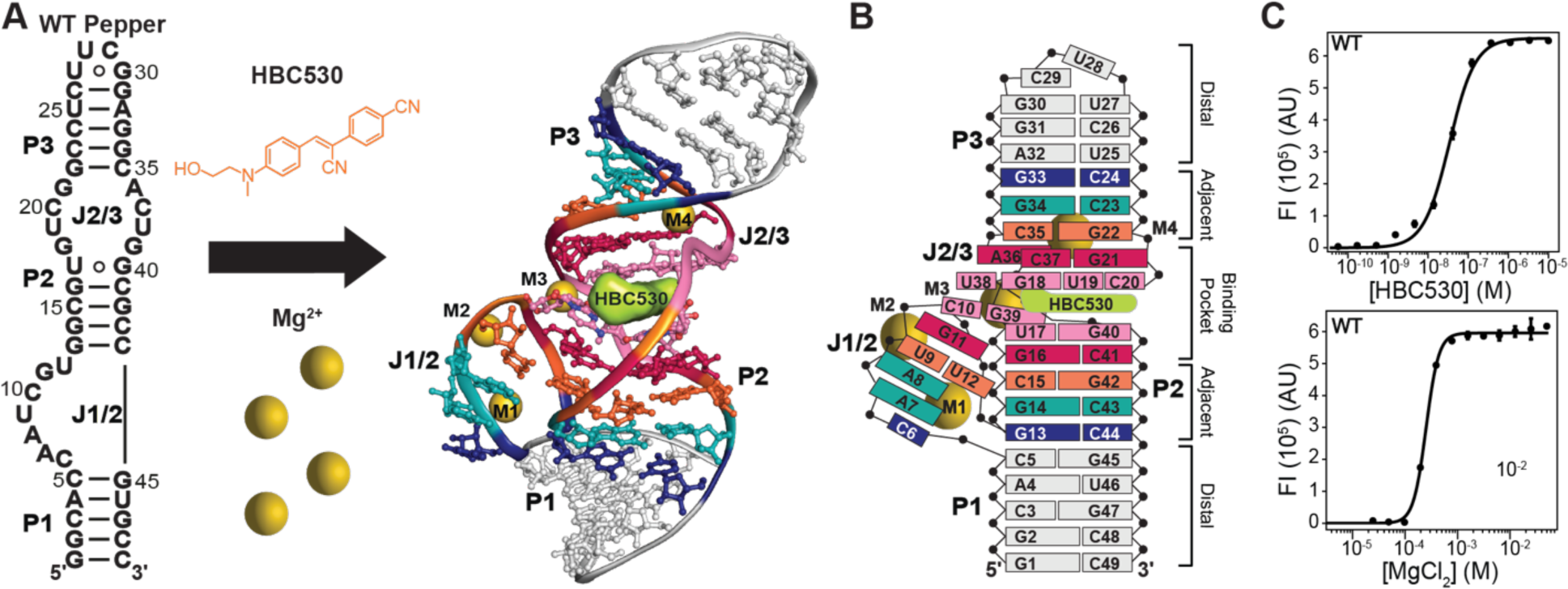
Pepper RNA aptamer binds HBC530. A) Schematic showing the secondary structure of WT Pepper aptamer, HBC530 and Mg^2+^ ligands, and a structural model adapted from the X-ray crystal structure (PDB ID 7EOH) and modified to match the sequence used in this study. Mg^2+^ cations are shown as yellow spheres and HBC530 in complex with RNA is shown in surface representation colored lime. RNA residues are colored by proximity to the HBC530 binding pocket where residues that directly interact with HBC530 are colored pink, residues in the binding pocket adjacent to direct residues are colored maroon, and residues adjacent to the binding pocket are colored orange, light blue, and dark blue from closest to farthest from the binding pocket. Residues considered distal to the binding pocket are colored gray. B) Schematic of RNA-ligand, RNA-RNA, and RNA-metal interactions. C) Representative plot of wild-type (WT) ligand binding assay for HBC530 (top) and MgCl_2_ (bottom) titrations.

Here, to gain quantitative insights into Pepper RNA-HBC530 complex formation and fluorescence activation we investigated the impact of Pepper RNA sequence on HBC530 recognition and Mg^2+^ coordination. In partnership with students in a Chemical Biology laboratory course, 53 Pepper RNA constructs were designed with point deletions, insertions, and/or substitutions. The apparent affinities to HBC530 and Mg^2+^, stoichiometries, and maximum fluorescence intensity (FI_max_) values were determined and compared to a wild-type (WT) Pepper RNA sequence. Consistent with previous studies, we found that substitutions to residues within the HBC530 binding pocket significantly reduced FI_max_ values. Moreover, the length and base-pair composition of stems bordering the HBC530 binding pocket, as well as the placement of the apical loop, impacted binding thermodynamics and fluorescence. A strong correlation was observed between Pepper affinity to HBC530 and Mg^2+^, in agreement with the established dependence of divalent cations on HBC530 recognition. Similarly, we found that the number of Mg^2+^ binding sites was correlated with both Mg^2+^ binding thermodynamics and FI_max_ values. In contrast, a modest correlation was observed between the binding affinity to HBC530 and FI_max_ values. We identified a class of Pepper constructs with near-WT affinity to HBC530 but significantly reduced FI_max_ values, indicating that RNA sequence plays a role in fluorescence activation that is decoupled from ligand binding affinity. Our results provide quantitative insights into the molecular mechanisms of Pepper-HBC530 recognition and fluorescence activation, revealing a complex interplay between binding thermodynamics and fluorescence.

## MATERIALS AND METHODS

### HBC530 synthesis

The HBC530 synthetic procedure was adapted from previous studies (12). The reaction scheme is provided in **Supplementary Fig. S1**. 0.2701 g (1.861 mmol) of 4-(cyanomethyl)benzonitrile and 20 mL of methanol were added to a 100 mL round-bottom flask with a stir bar. To this mixture, 0.5066 g (2.792 mmol) of 4-((2-hydroxyethyl)(methyl)amino)benzaldehyde was added. Once fully dissolved, 0.1 mL of piperidine was added to the reaction mixture under a nitrogen atmosphere. The temperature was maintained at 35-40 °C overnight with refluxing producing a dark orange solid. The mixture was concentrated using a rotary evaporator (Büchi), and the product was purified using flash column chromatography with 3% methanol in chloroform as the eluent. Fractions were pooled and concentrated using a rotary evaporator, yielding an orange solid weighing 0.504 g (1.656 mmol, 89% yield). Sample identity and purity was verified using Electrospray Ionization Mass Spectrometry (ESI-MS) and solution NMR spectroscopy (**Supplementary Fig. S2**). For ESI-MS, samples were dissolved in a mixture of water, acetonitrile, and 0.1% formic acid. For NMR, samples were dissolved in CDCl_3_ and ^1^H proton data was collected at 400 MHz.

### RNA construct design

The 49 nt wild-type (WT) construct used in this study was adapted from the previously reported Pepper aptamer sequence (12). Construct design was carried out in collaboration with a Chemical Biology teaching lab at the University of Nebraska - Lincoln over three student cohorts (N=36). Student cohorts consisted of advanced undergraduate and junior graduate students considered to be non-experts in RNA folding and structure. Students received two training sessions (totaling six hours) in RNA secondary structure prediction, RNA-ligand intermolecular interactions, literature reporting the impact of Pepper substitutions on fluorescence, and 3D structure visualization of a determined X-ray crystal structure of Pepper-HBC530 (PDB ID 7EOH) (13). With this introductory knowledge, students then designed a modified WT Pepper RNA construct with point substitutions, insertions, or deletions and used RNAfold (21,22) to maintain the hairpin secondary structure. These constructs were supplemented with additional point substitution constructs, as well as WT-like RNA constructs used in prior studies (12,13) as points of comparison for a total of 53 constructs (**Supplementary Table S1**). Constructs were named to denote the type of sequence variation, in which substitutions are indicated with WT residue number followed by the position and substituted nucleotide (e.g. C6U), deletions are indicated with a Δ symbol (e.g. ΔC6), base-pair substitutions are indicated with a hyphen (e.g. G16U-C41G), and insertions are indicated with .1N to indicate the addition of one nucleotide followed by the identity of the added nucleotide (e.g. C15.1C).

### RNA sample preparation

For all constructs (**Supplementary Table S1**), DNA templates were either purchased from Integrated DNA Technologies (IDT) or generated from primers designed using Primerize (23) and purchased from IDT. Polymerase chain reaction (PCR) overlap extension was performed to synthesize the DNA template with 20 cycles, 62-72 °C annealing temperature, and 5 μM each forward and reverse primers. The PCR product size was verified by agarose gel electrophoresis, followed by purification using a DNA clean and concentrator kit (Zymo Research). The DNA concentration was determined using Beer-Lambert’s law after measuring the absorbance of the samples with a Nanodrop spectrophotometer. RNA was transcribed at either 50 μL or 500 μL scale reactions in transcription buffer (40 mM Tris, pH 8.0, 1 mM spermine, 0.01% Triton X-100) with 150-250 nM DNA template, 20-30 mM MgCl_2_, 4 mM each rNTP, and T7 RNA polymerase. RNA was purified by gel excision from 20% polyacrylamide gel electrophoresis (PAGE) using UV shadowing. The RNA was extracted from the gel by incubating at ambient temperature in crush and soak buffer (300 mM sodium acetate, 1 mM EDTA, pH 5.22) for 24 hours. RNA was filtered using a Steriflip device to remove polyacrylamide and concentrated using a 10 kDa centrifugal filter (Amicon). The RNA was further purified using an RNA clean and concentrator kit (Zymo Research) to remove trace polyacrylamide. The purified RNA was then annealed by diluting the samples in ultrapure water to a concentration of 1-2 μM, heating at 95 °C for 4 minutes, followed by snap cooling on ice for one hour. The RNA was then buffer exchanged into storage buffer (20 mM HEPES, 50 mM KCl, pH 6.0). Prior to performing binding assays, the sample homogeneity of working RNA stocks was evaluated using 20% denaturing and 10% native analytical PAGE to confirm sample purity and monomeric folding, respectively.

### Binding assays

Binding assays were conducted at 296.15 K using black 96-well plates (Thermo Scientific, Catalog #12-566-23) with 10 nM RNA in a total volume of 100 μL per well. HBC530 titrations were performed in HBC530 titration buffer (10 mM MgCl_2_, 20 mM HEPES, 50 mM KCl, pH 6.0) with 0-10 μM HBC530. MgCl_2_ titrations were performed in storage buffer and 10 μM HBC530. Due to the wide range of Mg^2+^ midpoint ([Mg^2+^]_1/2_) values (0.090 – 29 mM), titrations of 0-10 mM MgCl_2_ were performed for constructs with [Mg^2+^]_1/2_ values less than 0.3 mM, and titrations of 0-50 mM MgCl_2_ were performed for constructs with [Mg^2+^]_1/2_ values greater than 0.3 mM. Prepared plates were incubated in the dark for one hour prior to data collection. Raw fluorescence data was collected on a BioTek Cytation 1 instrument with Gen5 software package with an excitation wavelength of 485 nm and emission wavelength of 528 nm. Each plate included a negative control, in duplicate, where RNA was omitted from the reaction. This control was used to apply baseline correction to the raw fluorescence data. For each construct, at least two independent replicates were performed, and each replicate included three technical replicates.

An in-house Python notebook was used to fit data to a modified Hill equation: 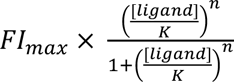 where FI_max_ is the maximum fluorescence intensity, [ligand] is the concentration of HBC530 or MgCl_2_, K is the apparent binding affinity (K_D_ or [Mg^2+^]_1/2_), and n is the Hill coefficient. The free energy of binding to HBC530 (1G_HBC530_) or Mg^2+^ (1G_Mg2+_) was calculated using Gibb’s free energy equation ΔG=RTln(K_D_ or [Mg^2+^]_1/2_) with a temperature of 296.15 K and a gas constant of 1.987 cal/mol•K. The mean fitted values and standard deviations are provided in **Supplemental Document 2**.

### Fluorimetry

Excitation and emission spectra were recorded on a Shimadzu RF-5301PC spectrofluorometer at ambient temperature. A saturated Pepper-HBC530 sample containing 1 μM RNA and 250 μM HBC530 was prepared in binding buffer (3 mM MgCl₂, 20 mM HEPES, 50 mM KCl, pH 6.0). For all samples, the excitation wavelength was 495 nm and the emission wavelength was collected from 500 nm to 800 nm. Data was collected at intervals of 0.2 nm, with excitation and emission bandwidths set to 3.0 nm. The instrument’s scan speed was set to slow, and sensitivity was set to high. Experiments were performed in triplicate.

### PDB survey of metal binding sites

Available X-ray crystal structures of Pepper-HBC complexes were downloaded from the PDB and aligned in PyMol. Structural models were manually inspected to identify reported metal ions in the X-ray crystal structural models. For the twelve available structures magnesium (all models), manganese (PDB ID 7EOI), sodium (PDB ID 7SZU), iridium hexammine (PDB ID 7EOG), and cesium (PDB ID 7EOJ) were observed. Iridium hexammine and cesium ions were excluded from further analysis. An in-house python script using the Bio.PDB package (24) in Biopython (version 1.85) (25) was generated to extract and group reported cations according to their coordinates. The RMSD was calculated for each group and was found to be 0.29-0.50 Å for the four most abundant metal sites. The code is available at https://github.com/eichhorn-lab/Pepper_variants.

## RESULTS

### Construct classification and global analysis of binding and fluorescence parameters

To systematically evaluate RNA Pepper sequence-structure-function relationships, we measured the apparent HBC530 binding affinity (K_D_) and maximum fluorescence intensity (FI_max_) values for 53 Pepper rationally designed constructs using established fluorescence-based HBC530 titration assays. In addition, we performed MgCl_2_ titration assays to measure the apparent Mg^2+^ midpoint ([Mg^2+^]_1/2_), or Mg^2+^ concentration in which one-half of the productive Pepper-HBC530 complex is formed. We classified constructs based on the proximity of point substitutions to the HBC530 binding pocket: ‘direct’ (contacting HBC530) and ‘indirect’ (neighboring) residues within the binding pocket, ‘adjacent’ (P2 stem, bottom half of P3 stem, and J1/2 residues that do not contribute to the binding pocket), and ‘distal’ (P1 stem, top half of P3 stem, apical loop) (**Fig. 1B**). This positional framework enables the identification of both local and distal contributions to Pepper-HBC530 recognition and fluorescence activation. Measured WT construct parameters are consistent with prior literature (13,14) showing nanomolar binding affinity and a stoichiometry of 1 HBC530 binding site per Pepper RNA. The near-identical FI_max_ values between HBC530 and MgCl_2_ titration experiments (R^2^ = 0.99, **Supplementary Fig. S3**) demonstrates excellent reproducibility in our experimental approach.

Diverse functional impacts were observed across the Pepper RNA construct library ranging from complete loss of fluorescence activation to better-than-WT performance. Eleven constructs showed little-to-no fluorescence response to HBC530 above baseline with high noise levels, and subsequently the data could not be fit (**Supplementary Document 2, Fig. S4**). When inspecting constructs with measurable binding (N=42) we observed a distribution spanning two orders of magnitude for FI_max_, K_D_, and [Mg^2+^]_1/2_ values (**Supplementary Fig. S5**). We next compared apparent binding affinities and observed 0.50-49-fold differences in K_D_ and 0.75-250-fold differences in [Mg^2+^]_1/2_ relative to WT (**Supplementary Fig. S6**), with differences in free energy (11G) values ranging from -0.1 kcal/mol to +3.4 kcal/mol (**Supplementary Fig. S7**).

To investigate the individual contributions of each junction in the binding pocket, we prepared constructs of the upper half (Pepper-top) and bottom half (Pepper-bottom) of Pepper, in addition to a truncated construct lacking the P3 stem (ΔP3) where J2/3 was converted into an apical loop. Pepper-bottom and ΔP3 constructs showed no detectable binding, confirming J2/3’s essential role in HBC530 recognition (**Supplementary Fig. S4**). In contrast Pepper-top retained binding; however, relative to WT the K_D_ was reduced 25-fold (11G_HBC530_ +1.89 ± 0.01 kcal/mol), [Mg^2+^]_1/2_ was reduced 88-fold (11G_Mg2+_ +2.63 ± 0.5 kcal/mol), and the relative FI_max_ was reduced to 8%. These results demonstrate that J2/3 is essential for HBC530 recognition while J1/2 has critical contributions to both binding thermodynamics and fluorescence.

### Binding pocket substitutions reveal complex structure-function relationships in HBC530 recognition and fluorescence

We first examined constructs with substitutions to residues that directly contact HBC530, which include C10-G39 to the side of HBC530, G18/U19/C20/U38 above HBC530, and U17•G40 underneath HBC530 (**Fig. 2A-C**). In addition to RNA-HBC530 contacts, the binding pocket contains numerous RNA-RNA hydrogen bonds: C10 (J1/2)-U17 (P2), C10 (J1/2)-G39 (J2/3), U17•G40 (P2), G18-U19-C20 (J2/3), U38-G39 (J2/3), and U19-U38 (**Fig. 2A-E**). All constructs with substitutions at positions that directly contact HBC530 had reduced FI_max_ values compared to WT, and several constructs showed little-to-no fluorescence activation (**Figs. 2F, S4**). For example, ΔC10 (J1/2) showed a 46-fold reduction in K_D_ (11G_HBC530_ +2.25 ± 0.01 kcal/mol) with a reduction in FI_max_ to 1.8% relative to WT (**Figs. 2F, S5**).

**Figure 2.**
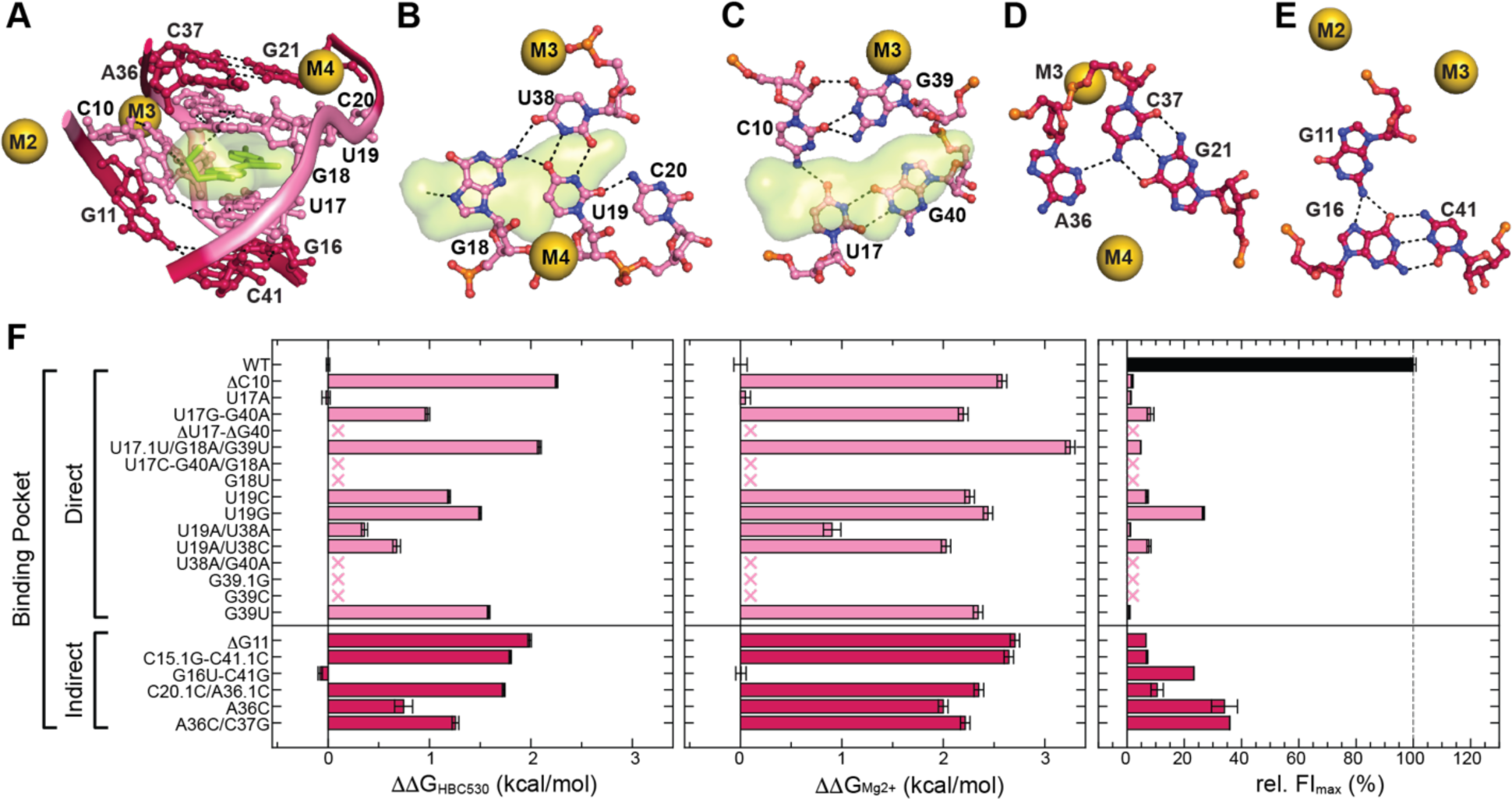
Mutational analysis of the HBC530 binding pocket reveals critical residues influencing binding thermodynamics and fluorescence intensity. A) Side view of the HBC530 binding pocket. The coloring scheme is the same as shown in Figure 1. B-C) Top view of residues that directly interact with HBC530 B) above and C) below HBC530. D-E) Top view of base triples D) above and E) below residues that directly contact HBC530. F) Bar plots for *left*: difference in free energy of binding HBC530 compared to WT (11G_HBC530_), *center*: 11G_Mg2+_, and *right*) relative fluorescence intensity maximum (FI_max_) values compared to WT Pepper RNA construct. Constructs showing no detectable fluorescence activation are marked with an x.

The U17•G40 wobble pair has been previously shown to be important for HBC530 recognition (12–14). As expected ΔU17•1G40, which also shortens the P2 stem by one base-pair, showed little fluorescence activation (**Supplementary Fig. S4**). Replacing the U•G wobble pair with an A•G sheared pair (U17A) did not impact K_D_ (11G_HBC530_ -0.02 ± 0.04 kcal/mol) but significantly reduced FI_max_ to 1.3% relative to WT. In contrast, a G•A sheared pair at this position (U17G•G40A) reduced the K_D_ 5-fold (11G_HBC530_ +0.98 ± 0.02 kcal/mol) and reduced FI_max_ to 8% relative to WT (**Supplementary Fig. S5**). These two constructs showed significantly different magnesium dependence, where U17A has near-WT values 11G_Mg2+_ +0.05 ± 0.05 kcal/mol) while U17G•G40A has 42-fold reduced [Mg^2+^]_1/2_ values (11G_Mg2+_ +2.2 ± 0.05 kcal/mol) (**Fig. 2F**). These results confirm the importance of the U17•G40 wobble pair on fluorescence activation.

Some substitutions likely impair HBC530 recognition through multiple mechanisms: blocking HBC530 access, removing direct RNA-HBC530 contacts, or disrupting the RNA-RNA interaction network. For example, G39C can form a G18-C39 base-pair that would occlude the HBC530 binding cavity (**Supplementary Fig. S8**) and must first be broken to enable a productive bound conformation, explaining the observed loss of fluorescence. In contrast G39U, which can form a G18•U39 wobble pair (**Supplementary Fig. S8**), retained binding to HBC530 but with a 15-fold reduction in K_D_ (11G_HBC530_ +1.58 ± 0.01 kcal/mol) and reduction in the relative FI_max_ to 1% (**Fig. 2F**). G18U, which can form a U18•G39 wobble pair (**Supplementary Fig. S8**) and also removes the hydrogen bond between the G18 N7 and HBC530 OH group, abolishes fluorescence activation. At the top of J2/3, the single point A36C substitution, which can form a G21-C36 base-pair that must unpair to form the G21-C37•A36 base triple in the HBC530-bound conformation (**Supplementary Fig. S8**), modestly reduced the K_D_ 4-fold (11G_HBC530_ +0.74 ± 0.09 kcal/mol). Further closing J2/3 by an additional base-pair (A36C/C37G) further reduced binding 8-fold (11G_HBC530_ +1.26 ± 0.03 kcal/mol). Despite observed differences in binding thermodynamics, these constructs showed nearly identical relative FI_max_ values (34% and 36%, respectively), suggesting that fluorescence is more sensitive than binding thermodynamics in this region. These differential outcomes underscore the intricate balance between competing intramolecular interactions and productive ligand binding.

Beyond direct RNA-HBC530 interactions, we investigated the impact of disrupting proximal structural elements within the binding pocket. Adjacent to residues that directly interact with HBC530 are G21-C37•A36 (J2/3) and G11•G16-C41 (J1/2 and P2) base-triples (**Fig. 2D-E**). Substitutions at these sites were overall less deleterious compared to residues that directly interact with HBC530, and no constructs abolished binding (**Fig. 2F**). Constructs with the greatest deleterious impact had deletions or insertions: ΔG11 (J1/2), C15.1G-C41.1C (P2), and C20.1C/A36.1C (J2/3) (**Fig. 2F**). Similar to U17•G40 variant constructs, substituting the G-C base-pair directly below U17•G40 with another G•U wobble pair (G16U•C41G) did not impact binding thermodynamics (11G_HBC530_ -0.08 ± 0.02 kcal/mol and 11G_Mg2+_ -0.005 ± 0.05 kcal/mol) but reduced fluorescence to ∼23% relative to WT (**Fig. 2F**). Collectively, these results reveal that while direct ligand contacts are essential for fluorescence, adjacent structural elements fine-tune both binding thermodynamics and fluorescence, with some constructs decoupling these two properties.

### P2 stem-J1/2 architecture and composition modulate binding thermodynamics and fluorescence

The five base-pair P2 stem serves as a structural scaffold that positions the J1/2 loop relative to J2/3, while J1/2 residues form both direct HBC530 contacts and tertiary interactions with J2/3 and the P2 major groove (**Fig. 3A-D**). To understand how this integrated architecture contributes to binding thermodynamics and fluorescence activation, we examined 16 constructs with modifications to either J1/2 or P2 (**Fig. 3E-F**). Shortening the P2 stem by one base-pair (ΔC15-ΔG42) reduced K_D_ 6-fold (11G_HBC530_ +1.06 ± 0.01 kcal/mol) and reduced the relative FI_max_ to 1%. When combined with a substitution of the adjacent G14-C43 base-pair to an A-U base-pair (G14A-C43U/ΔC15-ΔG42), fluorescence activation was abolished (**Supplementary Fig. S4**). Similarly, extending the P2 stem by one base-pair (C15.1G-C41.1C) reduced the K_D_ 21-fold (11G_HBC530_ +1.80 ± 0.01 kcal/mol) and reduced the relative FI_max_ to 7% (**Fig. 3E**). Substituting the terminal base-pair in P2 from G-C to G•U (C44U) improved the K_D_ 2-fold (11G_HBC530_ -0.33 ± 0.05 kcal/mol); however, the FI_max_ reduced to 49% relative to WT. Together, these results demonstrate the importance of both the length and composition of the P2 stem on both binding thermodynamics and fluorescence.

**Figure 3.**
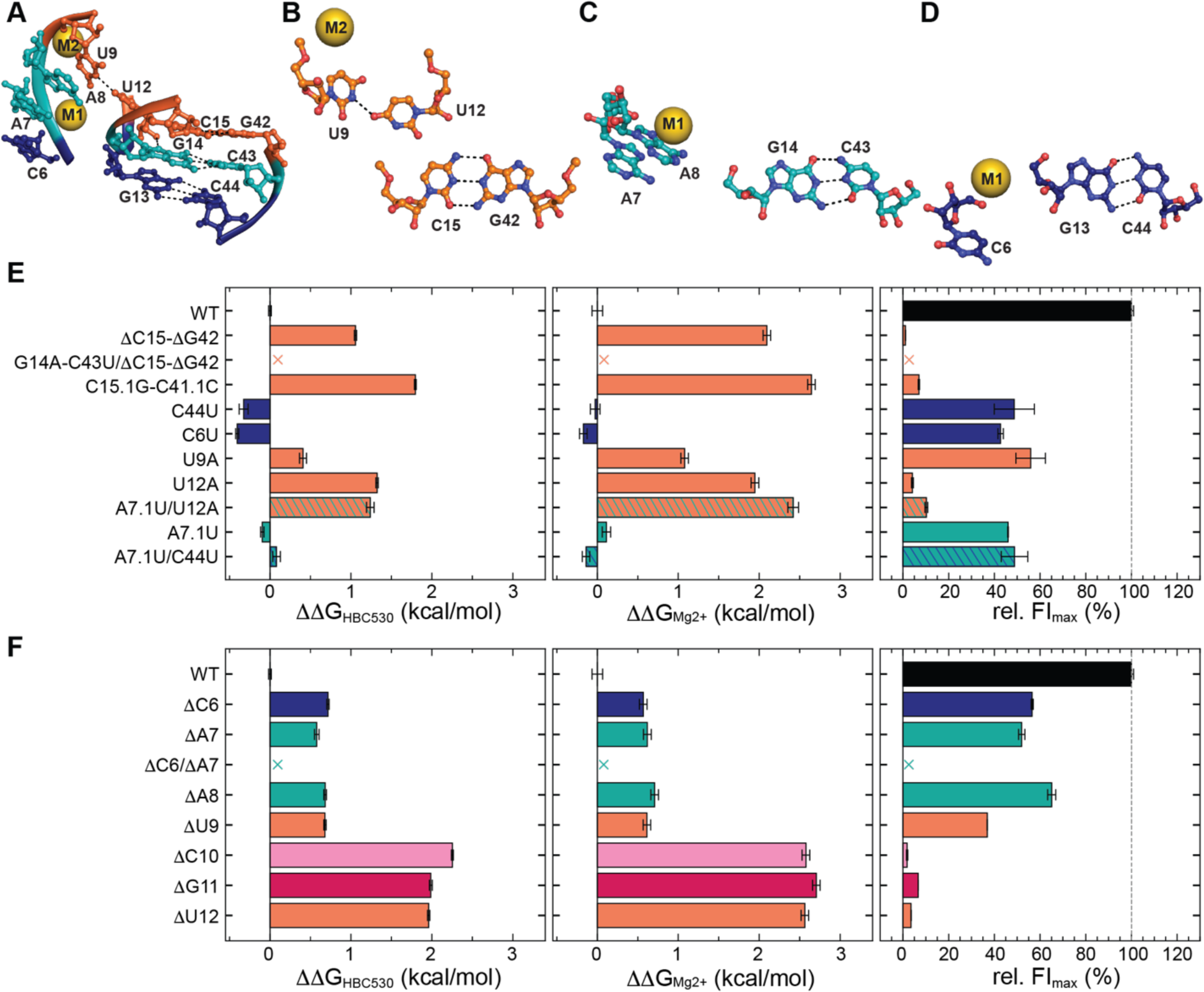
Substitutions to residues in the P2 stem and J1/2 loop adjacent to the HBC530 binding pocket have differential impacts on binding thermodynamics and fluorescence. A) Side view of the P2 stem region. The coloring scheme is the same as shown in Figure 1. B-D) Top view of residues B) below the HBC530 binding pocket, C) below residues in panel B, and D) below residues in panel C, at the bottom of the P2 stem. E-F) Bar plots of *left*: 11G_KD_, *center*: 11G_Mg2+_, and *right*) relative FI_max_ values for E) P2 and J1/2 substitutions and F) J1/2 point deletions. Constructs showing no detectable fluorescence activation are marked with an x.

We next probed the impact of residues in the J1/2 loop (**Fig. 3F**). Substituting C6, located at the base of J1/2, to a uridine (C6U) improved affinity 2-fold (11G_HBC530_ -0.40 ± 0.02 kcal/mol); however, the relative FI_max_ reduced to 43%. Below the binding pocket, U9•U12 (J1/2) docks in major groove of C15-G42 (P2) (**Fig. 3B-D**). Substitutions at these sites show differential impacts: U9A reduced K_D_ merely 2-fold (11G_HBC530_ +0.41 ± 0.04 kcal/mol) and retained 56% of FI_max_, while U12A reduced K_D_ nearly 10-fold (11G_HBC530_ +1.32 ± 0.01 kcal/mol) and FI_max_ was reduced to 4% relative to WT (**Fig. 3E**). A7 and A8 (J1/2) are positioned next to G14-C43 (P2) and are coordinated by the M1 Mg^2+^ cation (**Fig. 3A**). Inserting a uridine between A7 and A8 (A7.1U) had little impact on binding thermodynamics but reduced the relative FI_max_ to 46% (**Fig. 3E**). Systematic deletion of individual residues in J1/2 showed that while deletion of C6, A7, A8, or U9 was tolerated with modest reductions in binding thermodynamics (11G_HBC530_ ∼ +0.5 kcal/mol), deletion of C10, U11, or G12 showed severe impairment of both binding thermodynamics (11G_HBC530_ > +2 kcal/mol) and relative FI_max_ values (**Fig. 3F**). These results demonstrate the sequence and positional dependence of J1/2 residues in binding thermodynamics and fluorescence.

We next examined whether structural elements in J1/2 have cooperative interactions on binding thermodynamics. For example, RNA constructs were generated with either ΔC6, ΔA7, or ΔC6/ΔA7 in the J1/2 loop. While deletion of either C6 or A7 was tolerated, the combined C6/A7 deletion abolished fluorescence. In contrast, A7.1U showed a slight stabilizing effect (11G_HBC530_ - 0.09 ± 0.02 kcal/mol). U12A showed a significant destabilizing effect (11G_HBC530_ +1.32 ± 0.01 kcal/mol), as well as the 2-point substitution A7.1U/U12A (11G_HBC530_ +1.24 ± 0.05 kcal/mol). A comparison of individual and 2-point substitutions shows that these two sites are not thermodynamically coupled (11G_HBC530_,_diff_ +0.01 ± 0.05 kcal/mol). However, A7.1U is thermodynamically coupled to C44U (P2) (11G_HBC530,diff_ +0.5 ± 0.08 kcal/mol). Individual A7.1U and C44U substitutions are stabilizing (11G_HBC530_ -0.095 ± 0.02 and -0.33 ± 0.06 kcal/mol, respectively). However, the 2-point substitution A7.1U/C44U is destabilizing (11G_HBC530_ +0.08 ± 0.05 kcal/mol). Taken together, the Pepper RNA constructs with variations in the J1/2 loop and P2 stem show an important role in the length and composition of the P2 stem and residues C10-U12 in J1/2 in HBC530 recognition and fluorescence activation.

### Distal structural elements significantly influence binding thermodynamics and fluorescence

We next investigated whether more distal structural features also contribute to Pepper-HBC530 recognition and fluorescence. The original Pepper selection placed a UUCG tetraloop at the P3 stem (12), while subsequent X-ray crystallographic studies repositioned this loop to the P1 stem (13). To assess how tetraloop placement affects binding and fluorescence, we compared our WT construct with three circularly permutated variants containing the UUCG loop at the P1 stem (**Supplementary Fig. S9**). All three constructs showed increased FI_max_ values relative to WT (**Fig. 4A**). Maintaining the same sequence as WT but moving the UUCG apical loop to the P1 stem (WT-P1loop) showed near-identical binding thermodynamics and a 13% increase in FI_max_ (**Fig. 4A**). Swapping two base-pairs in the P3 stem (P3bpswap-P1loop) results in moderate 2-fold improvement in K_D_ (11G_HBC530_ -0.39 ± 0.03 kcal/mol) with a 20% increase in FI_max_ (**Fig. 4A**). The RNA sequence used in prior X-ray studies (X-ray), which differs from our WT construct with a G13A-C44U substitution and G-C base-pairs in P1 and P3 stems, also has 2-fold improved K_D_ (11G_HBC530_ -0.32 ± 0.02 kcal/mol) and 114% increased FI_max_ (**Fig. 4A**). These results show that increasing the thermodynamic stabilization of the P1 stem via tetraloop addition results in increased fluorescence.

**Figure 4.**
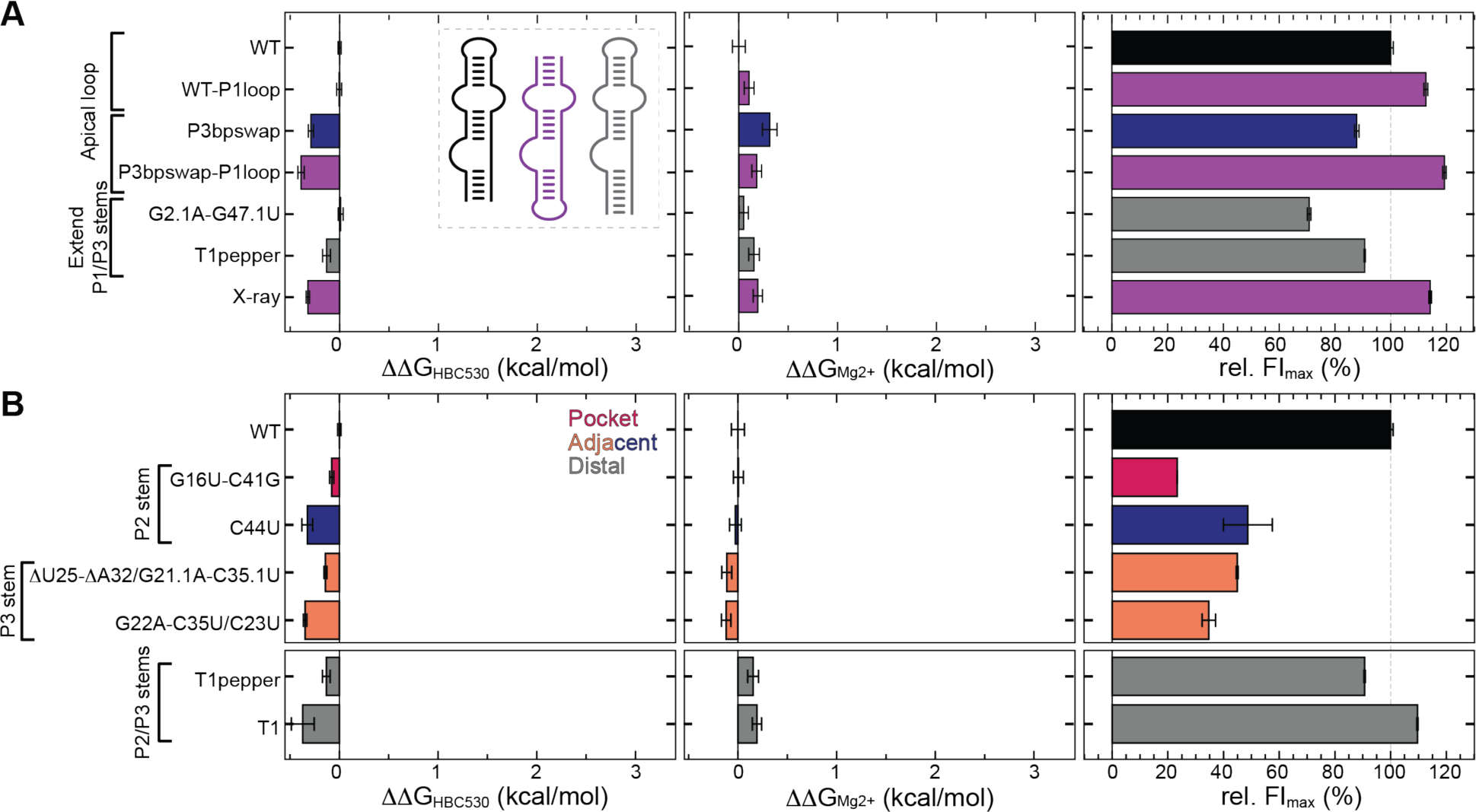
Base-pair substitutions outside the binding pocket influence both binding thermodynamics and fluorescence. The coloring scheme is the same as shown in Figure 1. A) WT-like constructs show increased fluorescence when placing the apical loop on the P1 stem. The inset cartoon schematic indicates apical loop placement, in which constructs colored purple have a P1-apical loop, and constructs colored gray have P1 and/or P3 stem extensions. B) Replacing G-C base pairs with G•U wobble pairs or A-U base-pairs in either P2 or P3 stems shows improved binding thermodynamic parameters but reduced relative FI_max_ values. The inset legend indicates the location of point substitutions. From left to right: Bar plots of *left*: 11G_KD_, *center*: 11G_Mg2+_, and *right*) relative FI_max_ values

To explore the impact of base-pair thermodynamic stability on Pepper-HBC530 binding thermodynamics and fluorescence, G-C base-pairs in the P2 or P3 stems were substituted with either G•U wobble pairs or A-U base-pairs, particularly at terminal base-pairs (**Fig. 4B**). Surprisingly, all constructs showed moderate favorable binding thermodynamics for both HBC530 and Mg^2+^ affinities (**Fig. 4B**). However, the relative FI_max_ values were reduced to 23-49%, suggesting that while reducing the thermodynamic stability of either the P2 or P3 stems results in favorable binding thermodynamics, fluorescence is impaired. The T1 construct is derived from the original Pepper RNA sequence from the selection while T1pepper is the T1 sequence with A13G-U44C/A22G substitutions (12) (**Supplementary Fig. S9**). These substitutions increase the thermodynamic stability of the terminal base-pairs of P2 and P3 stems by replacing A-U and A•C pairs, respectively, with G-C base-pairs. Similar to our previous observation T1 shows both a modest improvement in K_D_ and a 20% increased FI_max_ value relative to T1pepper (**Fig. 4B**). Together, these data show that the thermodynamic stability of each of the Pepper RNA stems have differential impacts on binding thermodynamics and fluorescence intensity.

### Ligand binding and fluorescence are decoupled functional properties

We next sought to identify global relationships between the measured parameters 11G_HBC530_, 11G_Mg2+_, and relative FI_max_. For constructs with measurable binding, we observed a modest correlation when comparing the relative FI_max_ to either the 11G_HBC530_ (**Fig. 5A**) or 11G_Mg2+_ (R^2^ = 0.49 and 0.45, respectively) (**Fig. 5B**). We identified several constructs with negligible or negative 11G_HBC530_ values, indicating favorable changes to the free energy of binding, but displayed a wide range of FI_max_ values from 1-120% compared to WT. For example, U17A has 1% relative FI_max_ and WT-P1loop has 120% relative FI_max_, but both exhibit near-WT binding thermodynamics. In contrast, a strong correlation was observed when comparing 11G_HBC530_ to the 11G_Mg2+_ (R^2^ = 0.88) (**Fig. 5C**) consistent with the importance of Mg^2+^ in HBC530 binding to the Pepper aptamer. These data show that reduced affinity to HBC530 is correlated with reduced affinity to Mg^2+^.

**Figure 5.**
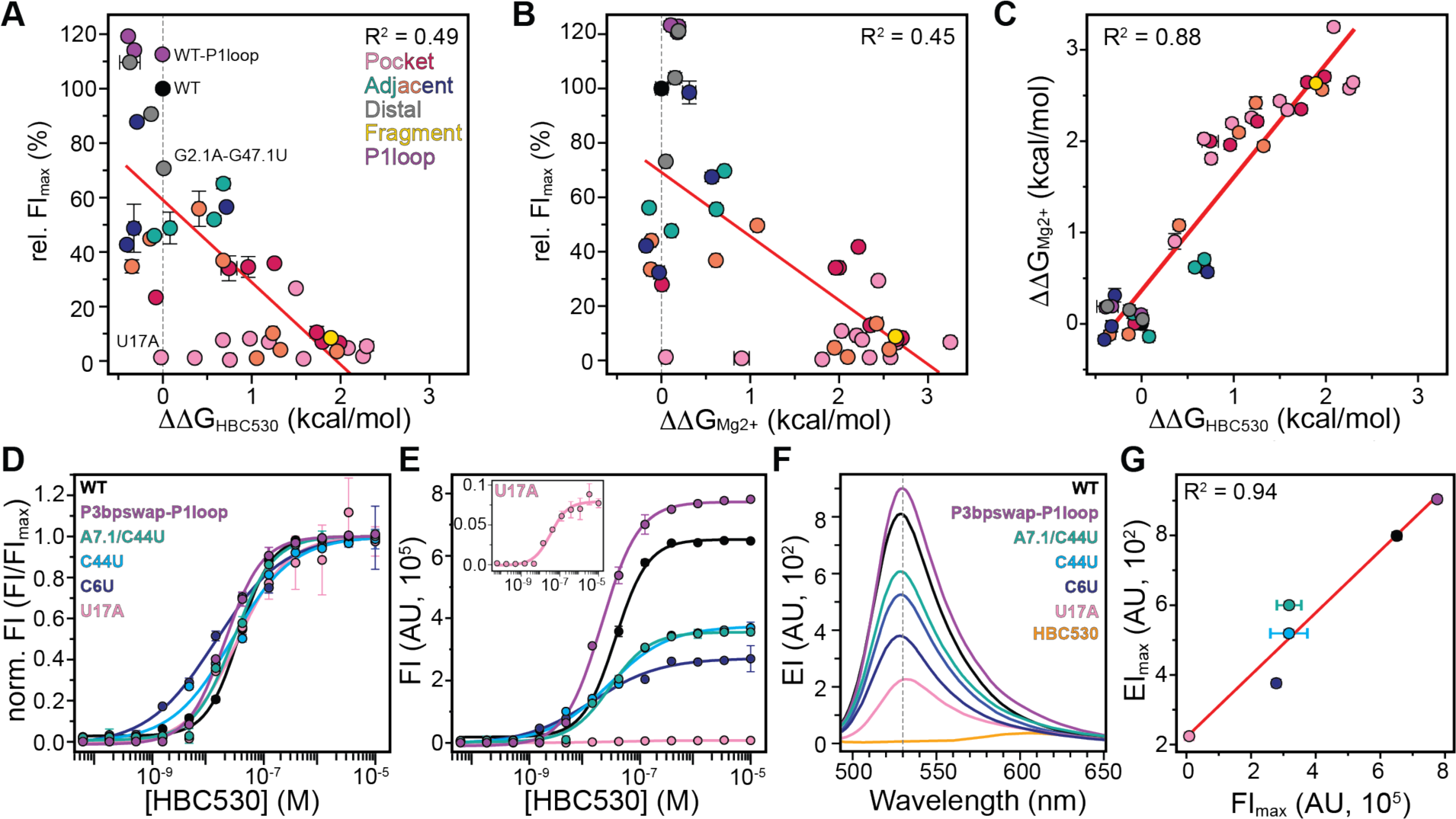
Identification of Pepper constructs with decoupled binding thermodynamics and fluorescence. A) Correlation plots of relative FI_max_ and 11G_HBC530_ and B) relative FI_max_ and 11G_Mg2+_ shows modest correlation between binding thermodynamics and fluorescence. Vertical dashed line indicates the WT 1G value (11G = 0), and the fit line is shown in red. C) Correlation plot of 11G_HBC530_ and 11G_Mg2+_ shows excellent agreement, D-E) Representative HBC530 titration data of selected constructs with near-WT affinity to HBC530 D) with or E) without normalized FI values, F) Fluorimeter measurement of emission intensity (EI), with the maximum emission wavelength indicated by a vertical dashed gray line, G) Correlation plot of the fluorimeter emission intensity maxima (EI_max_) values and HBC530 titration FI_max_ values show excellent agreement.

From these studies we identified 17 constructs with apparent K_D_ values within 2-fold of WT and selected a subset of constructs for additional analysis with substitutions to the HBC530 binding pocket (U17A), J1/2 (C6U) and adjacent P2 stem (C44U, A7.1U/C44U), and apical loop placement (P3bpswap-P1loop) (**Fig. 5D-G**). To rule out potential artifacts from the binding assays, we performed fluorimetry experiments using a saturated HBC530-bound complex for each RNA construct. All constructs showed the expected Stokes shift with emission wavelength maxima at 530 nm (**Fig. 5F**). However, we observed a range of maximum emission intensities for each construct (**Fig. 5F**) that have excellent agreement to FI_max_ values from the binding assays (R^2^ = 0.94) (**Fig. 5G**). From these experiments, we conclude that nucleotides in Pepper within the HBC530 binding pocket as well as distal regions have decoupled binding thermodynamics and fluorescence.

### Mg^2+^ stoichiometry influences binding thermodynamics and fluorescence

To investigate metal coordination in the Pepper-HBC complex, we surveyed the positions of reported divalent cations across available X-ray crystal structures and compared to the WT Mg^2+^ binding stoichiometry measured in this study. Currently twelve high-resolution X-ray crystal structures are available of Pepper RNA aptamers bound to an HBC derivative (13,14), containing between 2-9 divalent cations in each structure, with an average of 5.3 ± 2.2 sites (**Supplementary Fig. S10**). For these structures, divalent cations were reported at M1 and M3 sites for all twelve structures and at M2 and M4 sites in nine and ten structures, respectively (**Figs. 1A, 6A-B, S10**). All other reported metal binding sites were present in 33% or fewer of the X-ray crystal structures (**Supplementary Fig. S10**). Our binding assays of the WT construct showed a stoichiometry of 3 Mg^2+^ binding sites (**Supplementary Document 2**), which is in reasonable agreement with the observation of 2-4 cations in the structural data.

**Figure 6.**
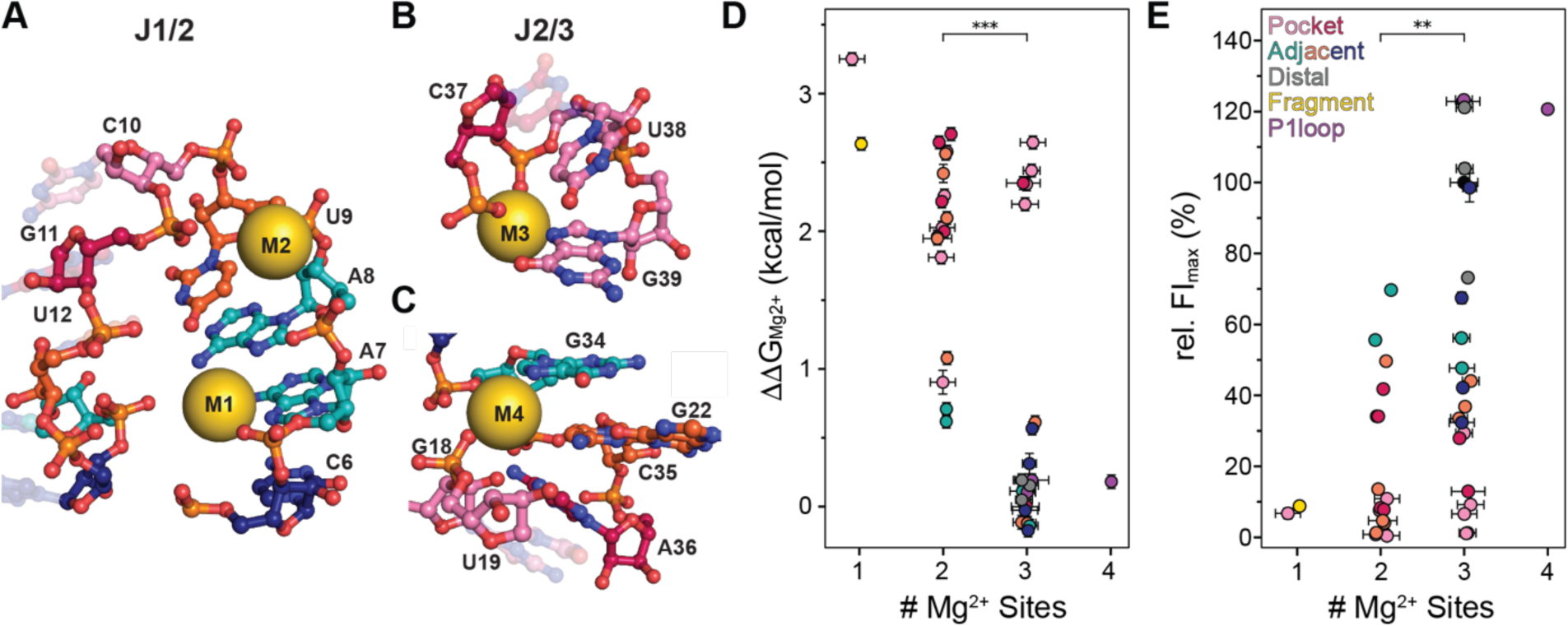
Mg^2+^ stoichiometry is correlated with binding thermodynamics and fluorescence intensity. The coloring scheme is the same as shown in Figure 1. A-C) Metal ion coordination in the Pepper-HBC complex. A) M1 and M2 bind to the J1/2 loop, B) M3 binds to the J2/3 loop, C) M4 binds to the J2/3 loop and bottom of the P3 stem. D) Comparison of 11G_Mg2+_ with the number of Mg^2+^ sites from MgCl_2_ titration assays shows that constructs with an increased number of Mg^2+^ sites have reduced 11G_Mg2+_ values (*** p-value 0.000030), E) Comparison of relative FI_max_ with the number of Mg^2+^ sites shows that constructs with an increased number of Mg^2+^ sites have increased FI_max_ values (** p-value 0.0024).

We next compared the Mg^2+^ binding stoichiometry across our construct library to either Mg^2+^ binding thermodynamics or fluorescence (**Figs. 6 and S11**). We inspected the stoichiometries of constructs with measurable binding and observed that most constructs fit to either 2 or 3 metal binding sites (17 and 22 constructs, respectively), with two constructs fitting to 1 metal binding site and one construct fitting to 4 sites (**Supplementary Document 2**). We observed a statistically significant decrease in 11G_Mg2+_ values for constructs with 3 sites compared to constructs with 2 sites (p-value 0.0024), indicating that increasing the number of metal binding sites results in more favorable binding thermodynamics (**Fig. 6D**). We observed a similar trend for the relative FI_max_ in which constructs with 3 sites have higher relative FI_max_ values compared to 2 sites (p-value 0.0030) (**Fig. 6E**). While 2-site constructs show a broad distribution of 11G_Mg2+_ values ranging between +0.5-3 kcal/mol, the 3-site constructs show a bimodal distribution where most constructs have near-WT Mg^2+^ binding thermodynamics (11G_Mg2+_ -0.1-+0.5 kcal/mol) (**Fig. 6D**). All five of the 3-site constructs with large 11G_Mg2+_ values contain substitutions in the HBC530 binding pocket (**Fig. 6D**). In summary, optimal magnesium coordination provides thermodynamic stabilization of the HBC530-bound complex conformation and subsequent fluorescence activation. Consequently, constructs with impaired metal binding at one or more sites exhibit both reduced binding thermodynamics and diminished fluorescence, further establishing metal binding as a critical determinant of Pepper RNA function.

## DISCUSSION

While the dependence of the Pepper aptamer sequence on fluorescence has been previously studied, the impact on ligand binding affinity, metal ion dependence, and thermodynamic contributions have not been comprehensively explored (12–14). Here, we surveyed 53 Pepper RNA constructs with insertions, deletions, and/or point substitutions to evaluate thermodynamic contributions to HBC530 recognition, Mg^2+^ binding, and fluorescence intensity. We found that, in general, substitutions within the binding pocket showed the most deleterious impacts on both binding thermodynamics and fluorescence. However, the overall correlation between binding thermodynamics and fluorescence was modest. Several constructs, including constructs with substitutions in the binding pocket, retained WT-like affinity to HBC530 but with significantly reduced FI_max_ values. Beyond the binding pocket, the length and composition of both the P2 stem and J1/2 loop were important for both binding thermodynamics and fluorescence. Similarly, we found that the base-pair composition of the P1 and P3 stems, and placement of the apical loop, impacted binding thermodynamics as well as fluorescence. Consistent with the important role of Mg^2+^ in Pepper-HBC recognition, we identified a strong correlation between HBC530 and Mg^2+^ binding thermodynamics and found that increasing the number of metal binding sites resulted in increased stabilization and higher fluorescence intensity. Importantly, we demonstrate that RNA novices can design functionally relevant RNA constructs that illuminate how aptamer sequence, structure, and thermodynamics intersect to influence aptamer-dye function.

Prior studies of Pepper RNA primarily focused on substitutions to the binding pocket, in particular C10 (J1/2) and residues in J2/3 (12–14). Consistent with previous studies, we find that substitutions to the U17•G40 wobble pair significantly reduce fluorescence (12–14). However, we identified that this site has decoupled fluorescence and binding thermodynamics. In addition, extending the P2 stem or substitutions to the terminal base-pair was previously shown to impact fluorescence (12–14). While we observe similar reductions to fluorescence, we find that reducing the thermodynamic stability of the terminal G13-C44 base-pair leads to overall favorable free energy of binding to HBC530. Likewise, the P1 and P3 stems distal to the binding pocket were previously reported to play structural roles and not impact fluorescence (18). Here, we show that the sequence composition of these stems subtly influences both binding thermodynamics and fluorescence, likely due to differences in the intrinsic RNA folding thermodynamics. Consistent with this model, increasing the thermodynamic stability of the P1 stem, whether by placing UUCG tetraloop (13,26) or adding a tRNA scaffold (20) resulted in increased fluorescence. These findings demonstrate that Pepper-HBC fluorescence enhancement is governed by a complex interplay between local inter- and intra-molecular interactions as well as global RNA stability.

Previous studies established that the affinity of the Pepper aptamer to HBC530 was dependent on Mg^2+^ (13). In this study, we quantified [Mg^2+^]_1/2_ values and compared the free energy of binding Mg^2+^ of constructs to WT. The [Mg^2+^]_1/2_ of a similar Pepper RNA-HBC530 construct was previously reported to be 1.2 mM with a stoichiometry of 1 (14). In our studies, the WT Pepper RNA construct has a 10-fold higher affinity to Mg^2+^ (0.11 mM) with a stoichiometry of 3. These differences are likely due to the reduced monovalent ionic strength in our study, as well as Pepper RNA sequence differences between the two studies (27). We found that constructs with reduced affinity to HBC530 also have reduced [Mg^2+^]_1/2_ values, indicating that additional Mg^2+^ is necessary to stabilize the Pepper-HBC530 tertiary structure and saturate HBC530 binding. These results demonstrate that Mg^2+^ binding and HBC530 recognition are thermodynamically coupled processes.

In this study, we showed site-specific decoupling of HBC530 binding and fluorescence in the Pepper RNA. A similar sequence dependence on fluorescence activation was observed for the Mango fluorogenic RNA aptamer (11), suggesting that this feature may be general to fluorogenic RNA aptamers. Importantly, our findings emphasize that binding alone is not sufficient for fluorescence activation, and that dual measurements of affinity and fluorescence are essential during aptamer selection. Further, when rationally designing substitutions to fluorogenic RNA aptamers the thermodynamic stability of distal elements should be considered. To design the next generation of fluorogenic aptamers, integrating structure-guided design alongside functional reporter assays during aptamer-dye pair development will be critical. A structure-guided approach was recently demonstrated to be successful in generating a novel fluorogen, named SALAD1, that binds to the Mango II aptamer with 3.5-fold higher fluorescence compared to TO1-biotin (28). In the future, research incorporating structure-guided design of RNA aptamers, alongside thermodynamic profiling of RNA and RNA-aptamer complexes, will further accelerate the rational engineering of aptamers with optimized fluorescence, stability, and *in vivo* functionality.

## DATA AVAILABILITY

Python scripts used to perform metal binding site analysis from available PDBs, and to analyze data and prepare figures are deposited in Github repository https://github.com/eichhorn-lab/Pepper_variants.

## SUPPLEMENTARY DATA

RNA construct sequences, fitted binding parameters, and additional supporting data are available as Supplemental Documents 1 and 2.

## AUTHOR CONTRIBUTIONS

CDE conceived and oversaw all aspects of the project. EKA prepared samples, performed data collection and analysis, and prepared figures. VJ prepared samples, and NTD prepared samples and performed initial data collection. PHD assisted with HBC530 synthesis and purification. CDE and EKA wrote the manuscript, and all authors provided feedback on the manuscript.

## ACKNOWLEDGEMENTS

We thank the students from the University of Nebraska – Lincoln CHEM437/837 course for RNA construct design and thoughtful discussions: Spring 2021: Ernest S. Atsrim, Momodou B. Camara, Korinne Erickson, Md Shirajur Rahman, Baba Yussif, and teaching assistant Gloricelly Roman Arocho; Spring 2023: Sachithra Nadeeshani Senanayake, Pinky Chowdhury, Sanduni Deenalattha, Shankar Ganesh, Nike Idowu, Serena Jentz, Xiangdong Liu, Shilla Owusu Ansah, Grace Palensky, Maggie Ramsay, and Harrison Wells; and Spring 2024: Oladeji Alabi, Oliver Brauning, Richard Buksch III, Addyson Calfee, Cristian Gonzalez, Sydney Goodman-Miller, Solenne Halman Gonzalez, Joshua Henning, Sakshi Jain, Laura Kirshenbaum, Yi Sheng Liong, Tharuki Malage, Claire Novak, Danushi Pathirage, Claire Weakly, and Lauren Wise. We thank Dr. Kurt Wulser for assistance collecting mass spectrometry data, and Dr. Martha Morton for supporting the NMR facility. We thank Dr. Joseph D. Yesselman and members of the Eichhorn lab for insightful discussions and feedback.

