## Supplementary Document 1 for "Pepper RNA variants reveal decoupling of HBC530 binding thermodynamics and fluorescence activation"

**List of supplementary information and figures:**

**Table S1.** Constructs used in this study.
**Figure S1.** Synthetic pathway for HBC530.
**Figure S2.** Characterization of HBC530 using NMR and ESI-MS.
**Figure S3**. Correlation plot of $FI_{max}$ from HBC530 and $MgCl_2$ titration experiments.
**Figure S4**. HBC530 titration experiments for constructs where binding could not be determined.
**Figure S5.** Bar plots of the fitted binding and fluorescence parameters for all constructs.
**Figure S6**. Bar plots of -fold changes relative to WT for all constructs.
**Figure S7**. Bar plots of the difference in free energy of binding for all constructs.
**Figure S8.** Secondary structures of selected Pepper RNA constructs showing substitutions at P2 and J2/3 junction that reduce the size of the J2/3 loop.
**Figure S9.** Secondary structures of Pepper RNA constructs that have an apical loop at the P1 stem, and constructs with base-pair substitutions at the P1, P2, or P3 stems.
**Figure S10.** Survey of available X-ray crystal structures of the Pepper-HBC complex.
**Figure S11**. Metal binding site analysis.

**Table S1.** Constructs used in the study

| Construct name | RNA construct sequence (5′–3′) | DNA construct sequence |
|---|---|---|
| *WT* | | |
| WT | GGCACCAAUCGUGGCGUGUCGG CCUCUUCGGAGGCACUGGCGCC GUGCC | CTAATACGACTCACTATAGGCACCAATCGTG GCGTGTCGGCCTCTTCGGAGGCACTGGCGC CGTGCC |
| *HBC530 Binding Pocket* | | |
| *Direct* | | |
| ΔC10 | GGCACCAAUGUGGCGUGUCGGC CUCUUCGGAGGCACUGGCGCCG UGCC | CTAATACGACTCACTATAGGCACCAATGTGG CGTGTCGGCCTCTTCGGAGGCACTGGCGCC GTGCC |
| U17A | GGCACCAAUCGUGGCGAGUCGGC CUCUUCGGAGGCACUGGCGCCG UGCC | TTCTAATACGACTCACTATAGGCACCAATCGT GGCGAGTCGGCCTCTTCGGAGGCACTGGCG CCGTGCC |
| U17G-G40A | GGCACCAAUCGUGGCGGGUCGG CCUCUUCGGAGGCACUGACGCCG UGCC | CTAATACGACTCACTATAGGCACCAATCGTG GCGGGTCGGCCTCTTCGGAGGCACTGACGC CGTGCC |
| ΔU17-ΔG40 | GGCACCAAUCGUGGCGGUCGGC CUCUUCGGAGGCACUGCGCCGU GCC | CTAATACGACTCACTATAGGCACCAATCGTG GCGGTCGGCCTCTTCGGAGGCACTGCGCCG TGCC |
| U17.1U/G18A/G39U | GGCACCAAUCGUGGCGUUAUCGG CCUCUUCGGAGGCACUUGCGCCG UGCC | CTAATACGACTCACTATAGGCACCAATCGTG GCGTTATCGGCCTCTTCGGAGGCACTTGCG CCGTGCC |
| U17C-G40A/G18A | GGCACCAAUCGUGGCGCAUCGGC CUCUUCGGAGGCACUGACGCCGU GCC | CTAATACGACTCACTATAGGCACCAATCGTG GCGCATCGGCCTCTTCGGAGGCACTGACGC CGTGCC |
| G18U | GGCACCAAUCGUGGCGUUUCGGC CUCUUCGGAGGCACUGGCGCCG UGCC | CTAATACGACTCACTATAGGCACCAATCGTG GCGTTTCGGCCTCTTCGGAGGCACTGGCGC CGTGCC |
| U19C | GGCACCAAUCGUGGCGUGCCGG CCUCUUCGGAGGCACUGGCGCC GUGCC | TTCTAATACGACTCACTATAGGCACCAATCGT GGCGTGCCGGCCTCTTCGGAGGCACTGGCG CCGTGCC |
| U19G | GGCACCAAUCGUGGCGUGGCGG CCUCUUCGGAGGCACUGGCGCC GUGCC | TTCTAATACGACTCACTATAGGCACCAATCGT GGCGTGGCGGCCTCTTCGGAGGCACTGGCG CCGTGCC |
| U19A/U38A | GGCACCAAUCGUGGCGUGACGGC CUCUUCGGAGGCACAGGCGCCGU GCC | CTAATACGACTCACTATAGGCACCAATCGTG GCGTGACGGCCTCTTCGGAGGCACAGGCGC CGTGCC |
| U19A/U38C | GGCACCAAUCGUGGCGUGACGGC CUCUUCGGAGGCACCGGCGCCG UGCC | CTAATACGACTCACTATAGGCACCAATCGTG GCGTGACGGCCTCTTCGGAGGCACCGGCGC CGTGCC |
| U38A/G40A | GGCACCAAUCGUGGCGUGUCGG CCUCUUCGGAGGCACAGACGCCG UGCC | CTAATACGACTCACTATAGGCACCAATCGTG GCGTGTCGGCCTCTTCGGAGGCACAGACGC CGTGCC |
| G39.1G | GGCACCAAUCGUGGCGUGUCGG CCUCUUCGGAGGCACUGGGCGC CGUGCC | CTAATACGACTCACTATAGGCACCAATCGTG GCGTGTCGGCCTCTTCGGAGGCACTGGGCG CCGTGCC |
| G39C | GGCACCAAUCGUGGCGUGUCGG CCUCUUCGGAGGCACUCGCGCCG UGCC | CTAATACGACTCACTATAGGCACCAATCGTG GCGTGTCGGCCTCTTCGGAGGCACTCGCGC CGTGCC |
| G39U | GGCACCAAUCGUGGCGUGUCGG CCUCUUCGGAGGCACUUGCGCCG UGCC | CTAATACGACTCACTATAGGCACCAATCGTG GCGTGTCGGCCTCTTCGGAGGCACTTGCGC CGTGC |
| *Indirect* | | |

| | | |
|---|---|---|
| ΔG11 | GGCACCAAUCUGGCGUGUCGGCCUCUUCGGAGGCACUGGCGCCGUGCC | TTCTAATACGACTCACTATAGGCACCAATCTGGCGTGTCGGCCTCTTCGGAGGCACTGGCGCCGTGCC |
| C15.1G-C41.1C | GGCACCAAUCGUGGCGGUGUCGGCCUCUUCGGAGGCACUGGCCGCCGUGCC | CTAATACGACTCACTATAGGCACCAATCGTGGCGGTGTCGGCCTCTTCGGAGGCACTGGCCGCCGTGCC |
| G16U-C41G | GGCACCAAUCGUGGCUUGUCGGCCUCUUCGGAGGCACUGGGGCCGUGCC | CTAATACGACTCACTATAGGCACCAATCGTGGCTTGTCGGCCTCTTCGGAGGCACTGGGGCCGTGCC |
| C20.1C/A36.1C | GGCACCAAUCGUGGCGUGUCCGGCCUCUUCGGAGGCACCUGGCGCCGUGCC | CTAATACGACTCACTATAGGCACCAATCGTGGCGTGTCCGGCCTCTTCGGAGGCACCTGGCGCCGTGCC |
| A36C | GGCACCAAUCGUGGCGUGUCGGCCUCUUCGGAGGCCCUGGCGCCGUGCC | CTAATACGACTCACTATAGGCACCAATCGTGGCGTGTCGGCCTCTTCGGAGGCCCTGGCGCCGTGCC |
| A36C/C37G | GGCACCAAUCGUGGCGUGUCGGCCUCUUCGGAGGCCGUGGCGCCGUGCC | CTAATACGACTCACTATAGGCACCAATCGTGGCGTGTCGGCCTCTTCGGAGGCCGTGGCGCCGTGCC |
| *Adjacent to the binding pocket* | | |
| ΔU9 | GGCACCAACGUGGCGUGUCGGCCUCUUCGGAGGCACUGGCGCCGUGCC | TTCTAATACGACTCACTATAGGCACCAACGTGGCGTGTCGGCCTCTTCGGAGGCACTGGCGCCGTGCC |
| U9A | GGCACCAAACGUGGCGUGUCGGCCUCUUCGGAGGCACUGGCGCCGUGCC | CTAATACGACTCACTATAGGCACCAAACGTGGCGTGTCGGCCTCTTCGGAGGCACTGGCGCCGTGCC |
| ΔU12 | GGCACCAAUCGGGCGUGUCGGCCUCUUCGGAGGCACUGGCGCCGUGCC | TTCTAATACGACTCACTATAGGCACCAATCGGGCGTGTCGGCCTCTTCGGAGGCACTGGCGCCGTGCC |
| U12A | GGCACCAAUCGAGGCGUGUCGGCCUCUUCGGAGGCACUGGCGCCGUGCC | CTAATACGACTCACTATAGGCACCAATCGAGGCGTGTCGGCCTCTTCGGAGGCACTGGCGCCGTGCC |
| ΔU25-ΔA32/G21.1A-C35.1U | GGCACCAAUCGUGGCGUGUCGAGCCCUUCGGGGCUACUGGCGCCGUGCC | CTAATACGACTCACTATAGGCACCAATCGTGGCGTGTCGAGCCCTTCGGGGCTACTGGCGCCGTGCC |
| ΔC15-ΔG42 | GGCACCAAUCGUGGGUGUCGGCCUCUUCGGAGGCACUGGCCCGUGCC | CTAATACGACTCACTATAGGCACCAATCGTGGGTGTCGGCCTCTTCGGAGGCACTGGCCCGTGCC |
| ΔA7 | GGCACCAUCGUGGCGUGUCGGCCUCUUCGGAGGCACUGGCGCCGUGCC | CTAATACGACTCACTATAGGCACCATCGTGGCGTGTCGGCCTCTTCGGAGGCACTGGCGCCGTGCC |
| A7.1U | GGCACCAUAUCGUGGCGUGUCGGCCUCUUCGGAGGCACUGGCGCCGUGCC | TTCTAATACGACTCACTATAGGCACCATATCGTGGCGTGTCGGCCTCTTCGGAGGCACTGGCGCCGTGCC |
| ΔA8 | GGCACCAUCGUGGCGUGUCGGCCUCUUCGGAGGCACUGGCGCCGUGCC | TTCTAATACGACTCACTATAGGCACCATCGTGGCGTGTCGGCCTCTTCGGAGGCACTGGCGCCGTGCC |
| ΔC6 | GGCACAAUCGUGGCGUGUCGGCCUCUUCGGAGGCACUGGCGCCGUGCC | TTCTAATACGACTCACTATAGGCACAATCGTGGCGTGTCGGCCTCTTCGGAGGCACTGGCGCCGTGCC |
| C6U | GGCACUAAUCGUGGCGUGUCGGCCUCUUCGGAGGCACUGGCGCCGUGCC | CTAATACGACTCACTATAGGCACTAATCGTGGCGTGTCGGCCTCTTCGGAGGCACTGGCGCCGTGCC |
| C44U | GGCACCAAUCGUGGCGUGUCGGCCUCUUCGGAGGCACUGGCGCUGUGCC | CTAATACGACTCACTATAGGCACCAATCGTGGCGTGTCGGCCTCTTCGGAGGCACTGGCGCTGTGCC |

| | | |
|---|---|---|
| *Multiple regions* | | |
| C20G/C37G | GGCACCAAUCGUGGCGUGUGGGCCUCUUCGGAGGCAGUGGCGCCGUGCC | CTAATACGACTCACTATAGGCACCAATCGTGGCGTGTGGGCCTCTTCGGAGGCAGTGGCGCCGTGCC |
| C10G/G11U/U12A | GGCACCAAUGUAGGCGUGUCGGCCUCUUCGGAGGCACUGGCGCCGUGCC | CTAATACGACTCACTATAGGCACCAATGTAGGCGTGTCGGCCTCTTCGGAGGCACTGGCGCCGTGCC |
| U9C/G11C/ΔC3-ΔG47 | GGACCAACCCUGGCGUGUCGGCCUCUUCGGAGGCACUGGCGCCGUCC | CTAATACGACTCACTATAGGACCAACCCTGGCGTGTCGGCCTCTTCGGAGGCACTGGCGCCGTCC |
| U12.1U/G14C-C43G/C15G-G42C | GGCACCAAUCGUUGCGGUGUCGGCCUCUUCGGAGGCACUGGCCGCGUGCC | CTAATACGACTCACTATAGGCACCAATCGTTGCGGTGTCGGCCTCTTCGGAGGCACTGGCCGCGTGCC |
| A7.1U/U12A | GGCACCUAAUCGAGGCGUGUCGGCCUCUUCGGAGGCACUGGCGCCGUGCC | CTAATACGACTCACTATAGGCACCTAATCGAGGCGTGTCGGCCTCTTCGGAGGCACTGGCGCCGTGCC |
| G14A-C43U/ΔC15-ΔG42 | GGCACCAAUCGUGAGUGUCGGCCUCUUCGGAGGCACUGGCUCGUGCC | CTAATACGACTCACTATAGGCACCAATCGTGAGTGTCGGCCTCTTCGGAGGCACTGGCTCGTGCC |
| G22A-C35U/C23U | GGCACCAAUCGUGGCGUGUCGAUCUCUUCGGAGGUACUGGCGCCGUGCC | CTAATACGACTCACTATAGGCACCAATCGTGGCGTGTCGATCTCTTCGGAGGTACTGGCGCCGTGCC |
| ΔC6/ΔA7 | GGCACAUCGUGGCGUGUCGGCCUCUUCGGAGGCACUGGCGCCGUGCC | CTAATACGACTCACTATAGGCACATCGTGGCGTGTCGGCCTCTTCGGAGGCACTGGCGCCGTGCC |
| A7.1U/C44U | GGCACCUAAUCGUGGCGUGUCGGCCUCUUCGGAGGCACUGGCGCUGUGCC | CTAATACGACTCACTATAGGCACCTAATCGTGGCGTGTCGGCCTCTTCGGAGGCACTGGCGCTGTGCC |
| P3bpswap | GGCACCAAUCGUGGCGUGUCGGCUCCUUCGGGAGCACUGGCGCCGUGCC | TTCTAATACGACTCACTATAGGCACCAATCGTGGCGTGTCGGCTCCTTCGGGAGCACTGGCGCCGTGCC |
| *Distal to HBC530 binding pocket* | | |
| T1pepper | GGACACCAAUCGUGGCGUGUCGGCCUGCUUCGGCAGGCACUGGCGCCGUGUCC | CTAATACGACTCACTATAGGGACACCAATCGTGGCGTGTCGGCCTGCTTCGGCAGGCACTGGCGCCGTGTCC |
| T1 | GGACACCAAUCGUAGCGUGUCGACCUGCUUCGGCAGGCACUGGCGCUGUGUCC | CTAATACGACTCACTATAGGGACACCAATCGTAGCGTGTCGACCTGCTTCGGCAGGCACTGGCGCTGTGTCC |
| G2.1A-G47.1U | GGACACCAAUCGUGGCGUGUCGGCCUCUUCGGAGGCACUGGCGCCGUGUCC | CTAATACGACTCACTATAGGACACCAATCGTGGCGTGTCGGCCTCTTCGGAGGCACTGGCGCCGTGTCC |
| *Fragment constructs* | | |
| ΔP3 | GGCACCAAUCGUGGCGUGUCGUACUGGCGCCGUGCC | CTAATACGACTCACTATAGGCACCAATCGTGGCGTGTCGTACTGGCGCCGTGCC |
| Pepper-top | GGGCGUGUCGGCCUCUUCGGAGGCACUGGCGCC | CTAATACGACTCACTATAGGGCGTGTCGGCCTCTTCGGAGGCACTGGCGCC |
| Pepper-bottom | GGGCACCAAUCGUGGCGCUUCGGGCGCCGUGCC | CTAATACGACTCACTATAGGGCACCAATCGTGGCGCTTCGGGCGCCGTGCC |
| *Apical loop on P1 stem* | | |
| P3bpswap-P1loop | GGAGCACUGGCGCCGUGCCUUCGGGCACCAAUCGUGGCGUGUCGGCUCC | CTAATACGACTCACTATAGGAGCACTGGCGCCGTGCCTTCGGGCACCAATCGTGGCGTGTCGGCTCC |

| | | |
|---|---|---|
| Xray | GGCGCACUGGCGCUGCGCCUUC GGGCGCCAAUCGUAGCGUGUCG GCGCC | CTAATACGACTCACTATAGGCGCACTGGCGC TGCGCCTTCGGGCGCCAATCGTAGCGTGTC GGCGCC |
| WT-P1loop | GAGGCACUGGCGCCGUGCCUUC GGGCACCAAUCGUGGCGUGUCG GCCUC | TTCTAATACGACTCACTATAGAGGCACTGGC GCCGTGCCTTCGGGCACCAATCGTGGCGTG TCGGCCTC |

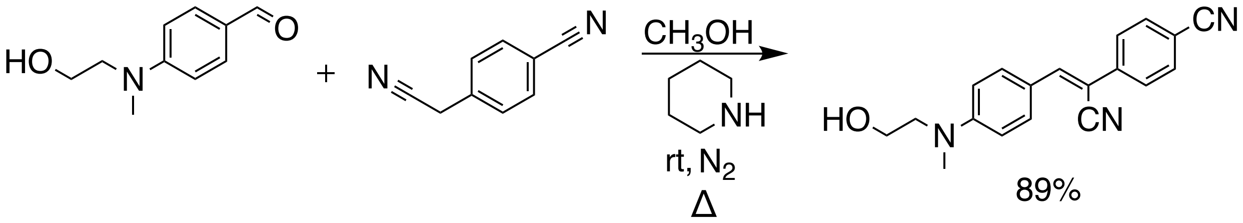


**Figure S1.** Synthetic pathway for HBC530.

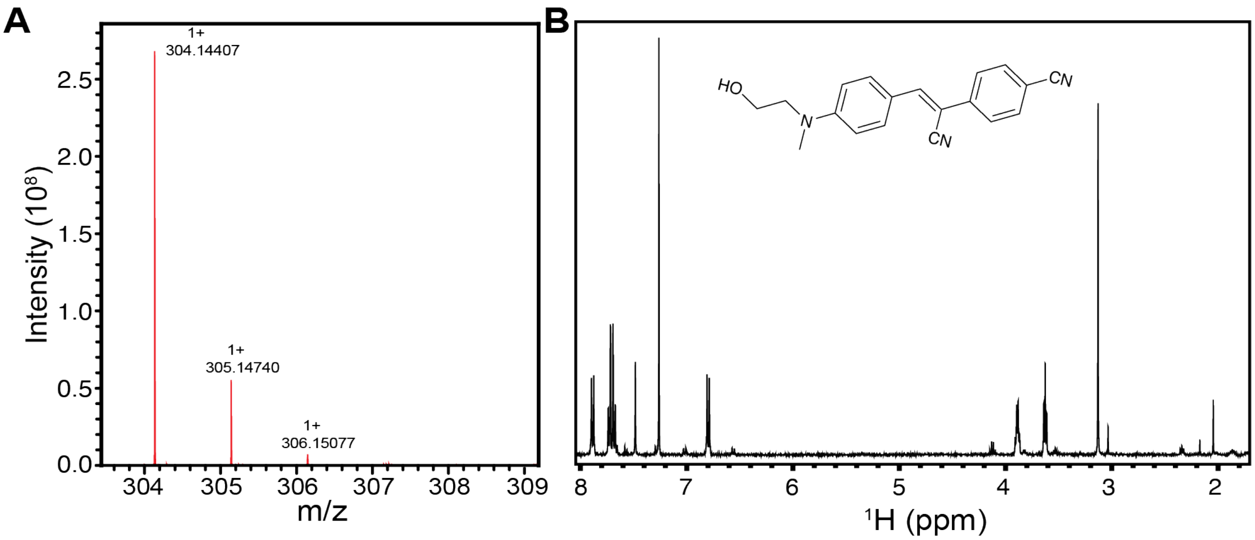


**Figure S2.** Characterization of HBC530 using ESI-MS and NMR. A) ESI-TOF mass spectrum of HBC530. Theoretical calculation: $C_{19}H_{18}N_3O$ [M+H]+: 304.1450, experimental: 304.14407. B) NMR data for HBC530 $^1$H NMR (400 MHz, $CDCl_3$) δ 7.89 (d, 2H), δ 7.70 (d, 4H), δ 7.48 (s, 1H), δ 6.80 (d, 2H), δ 3.89 (d, 2H), δ 3.63 (t, 3H), δ 3.13 (s, 3H)

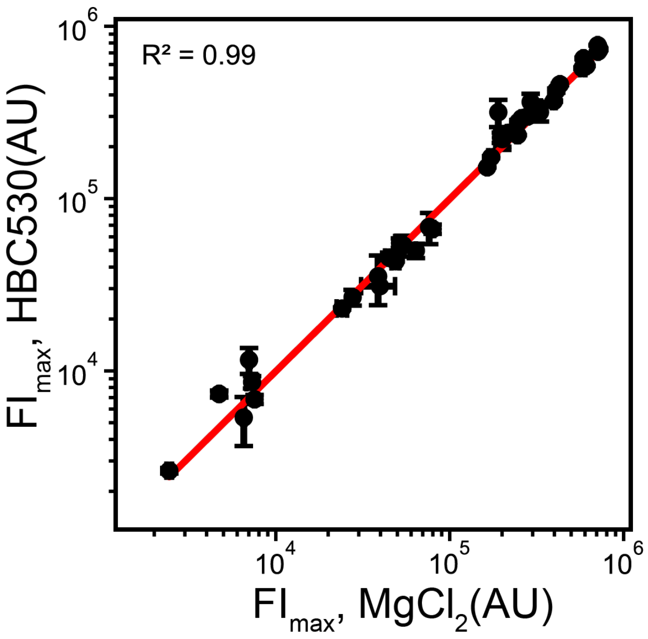


**Figure S3**. Correlation plot of $FI_{max}$ values from HBC530 and $MgCl_2$ titration experiments. AU indicates arbitrary units. The fit line is shown in red.

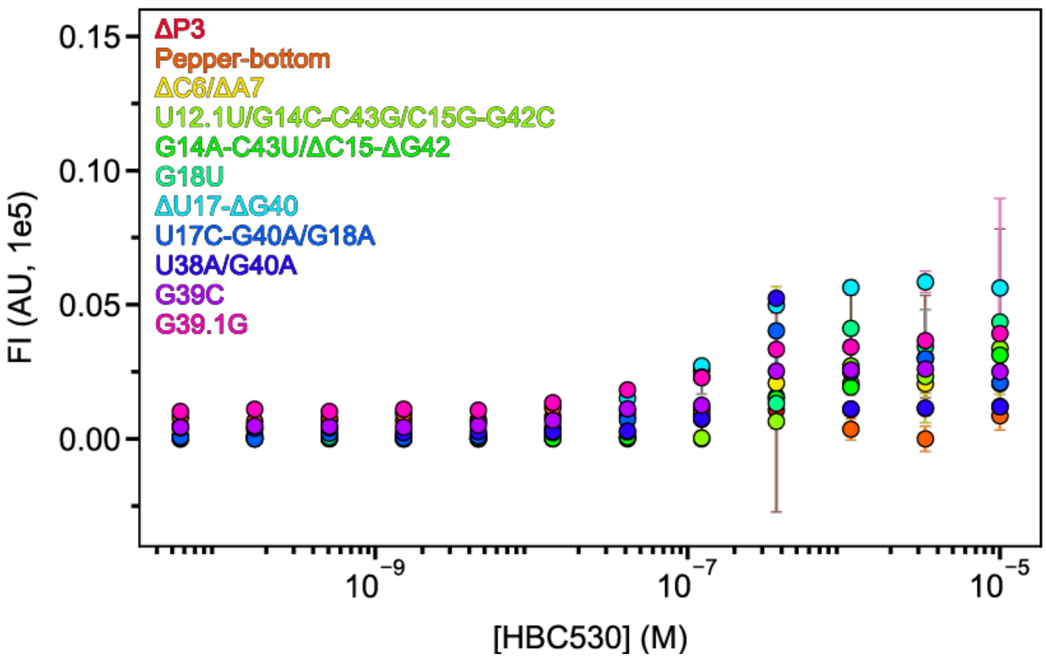


**Figure S4**. HBC530 titration experiments for constructs where binding could not be determined. Note that the WT Pepper-HBC530 complex has a maximum FI value of 6.5 x $10^5$ AU.

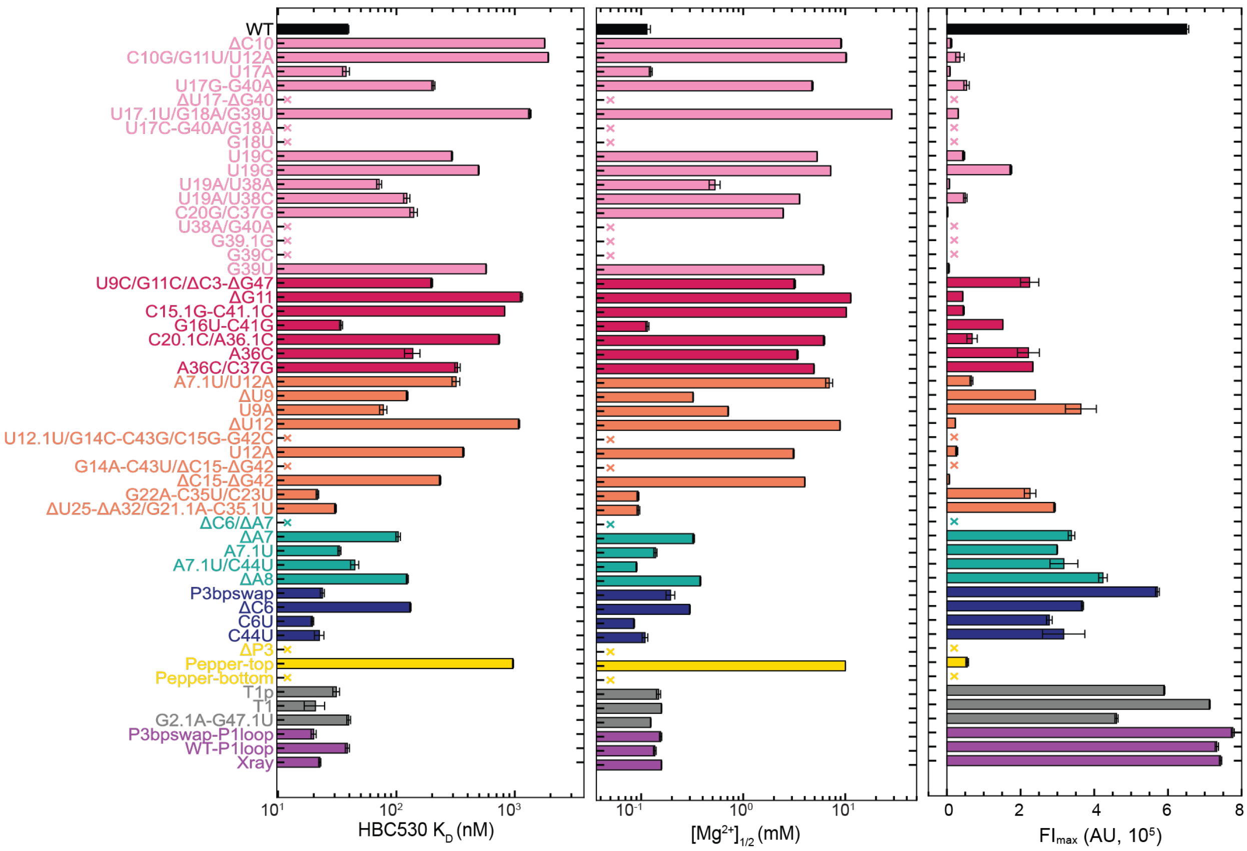


**Figure S5**. Bar plots of the fitted binding and fluorescence parameters for all constructs of (from left to right): the apparent HBC530 affinity, $Mg^{2+}$ midpoint, and maximum fluorescence intensity. X denotes constructs with low fluorescence, and fluorescence-based assays could not determine binding parameters.

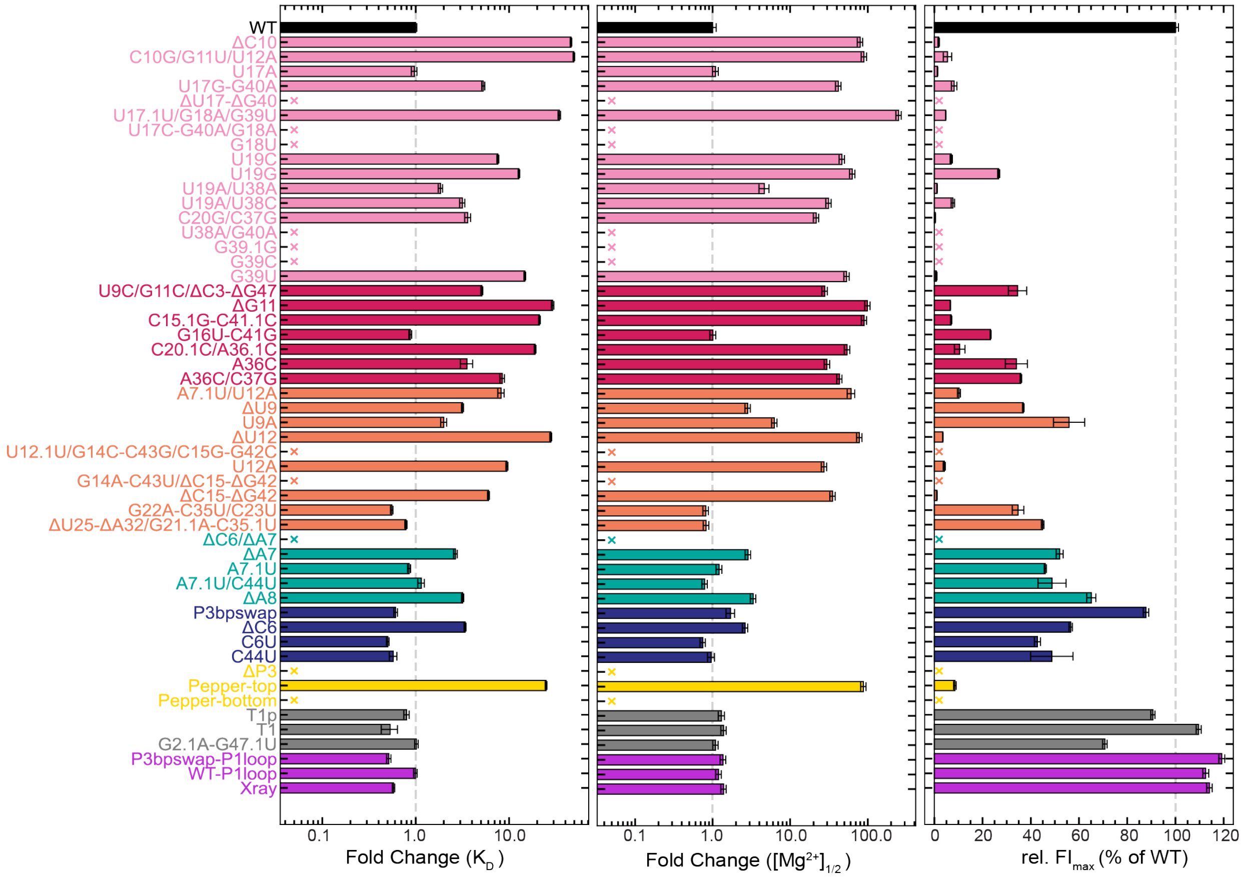


**Figure S6**. Bar plots of -fold changes relative to WT for all constructs. From left to right: HBC530 affinity fold change, $Mg^{2+}$ affinity fold change, and relative fluorescence intensity compared to WT. Fluorescence intensity data is shown from HBC530 titration experiments.

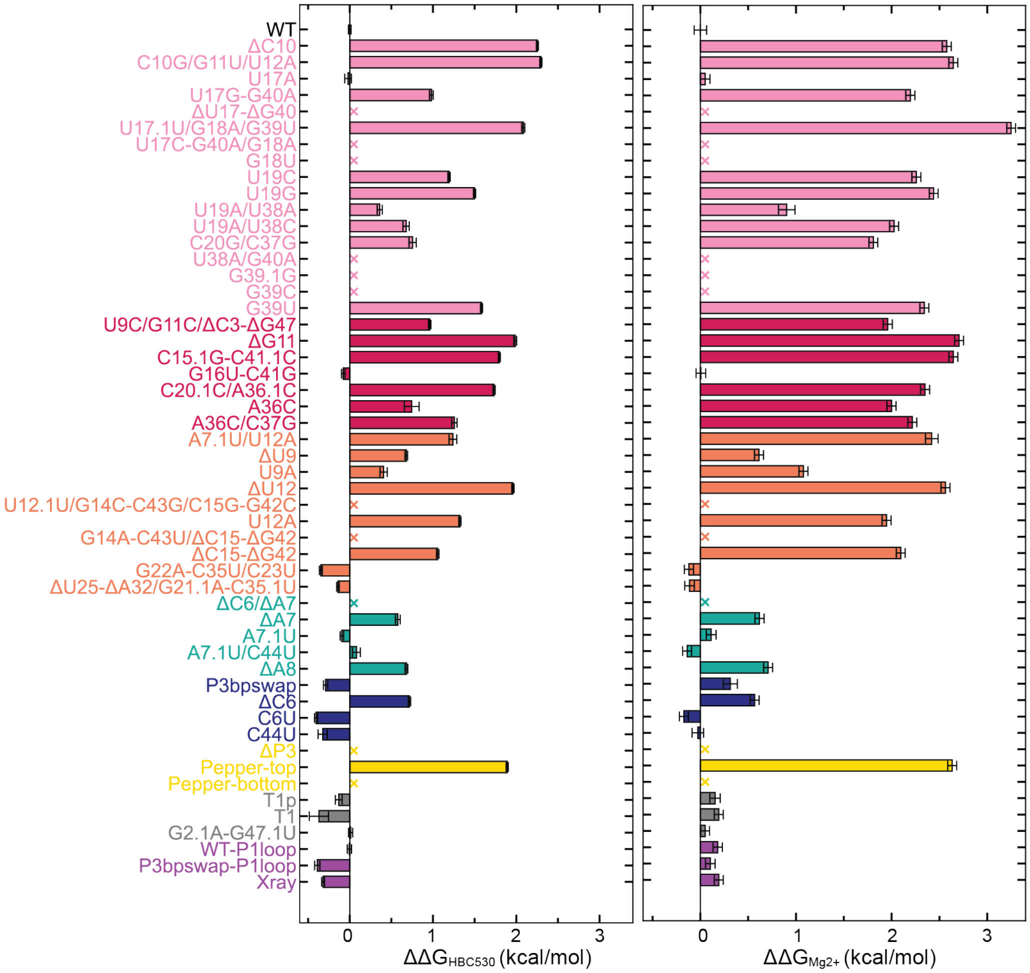


**Figure S7**. Bar plots of the difference in free energy of binding for all constructs.

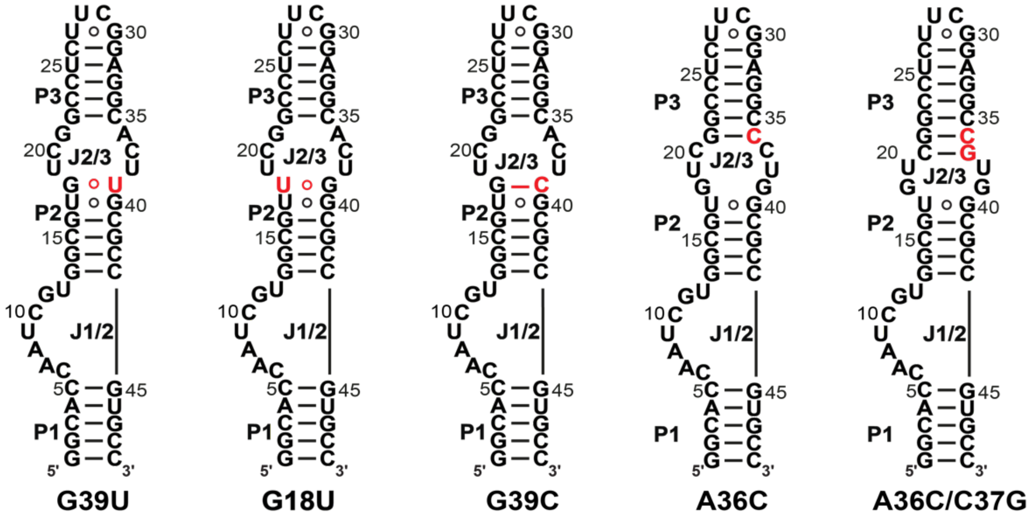


**Figure S8.** Secondary structures of selected Pepper RNA constructs showing substitutions at P2 and J2/3 junction that reduce the size of the J2/3 loop. Residues that are colored red indicate sequence differences compared to WT.

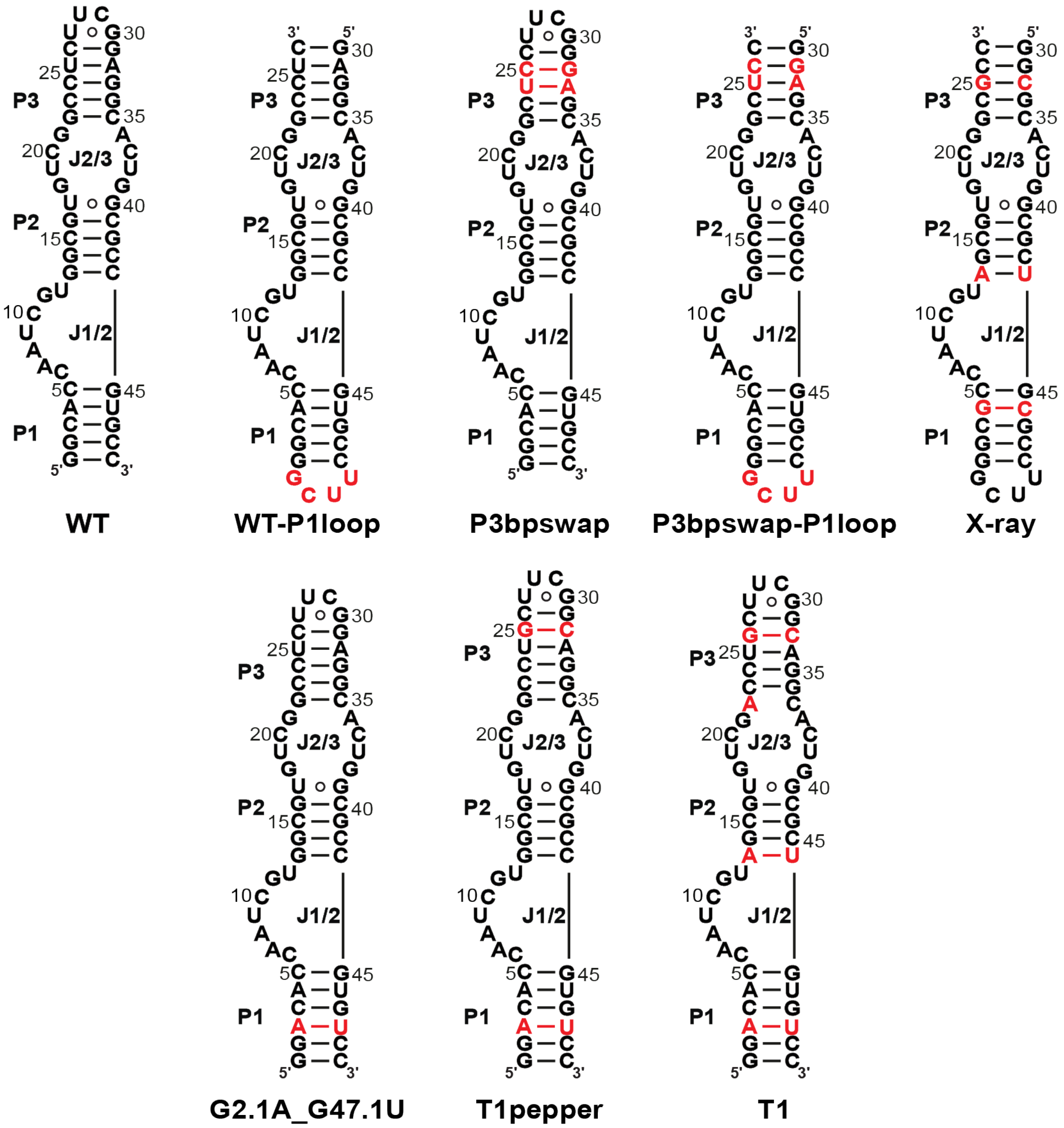


**Figure S9.** Secondary structures of Pepper RNA constructs that have an apical loop at the P1 stem, and constructs with base-pair substitutions at the P1, P2, or P3 stems. Residues that are colored red indicate sequence differences compared to WT.

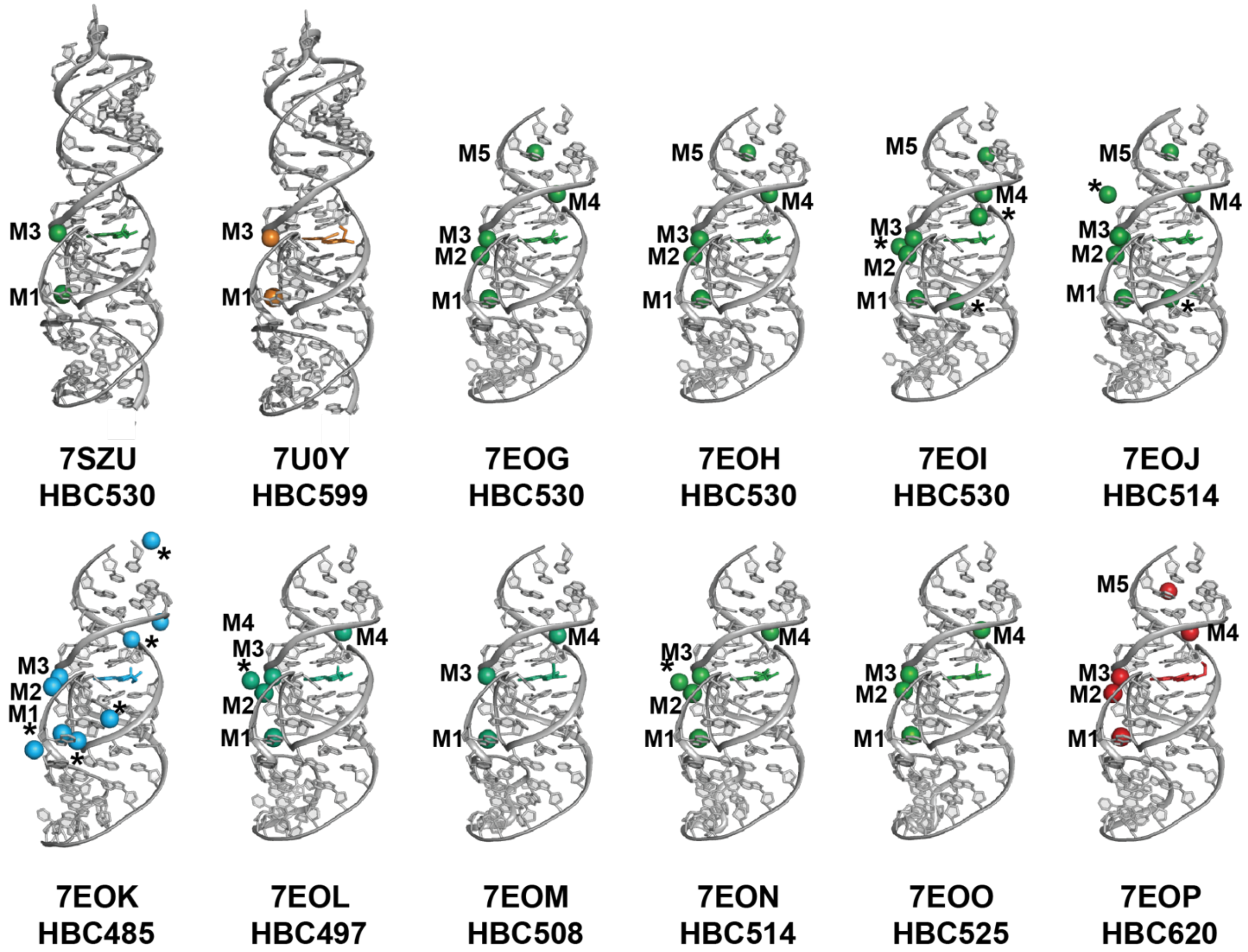


**Figure S10.** Survey of available X-ray crystal structures of the Pepper-HBC complex. The PDB ID and HBC derivative is labeled below each crystal structure. RNA is colored gray and HBC ligand and divalent cations are colored according to the HBC maximum emission wavelength. The divalent cations that are marked with an asterisk are additional metal ions not present in the Pepper-HBC530 complex structure (PDB ID 7EOH).

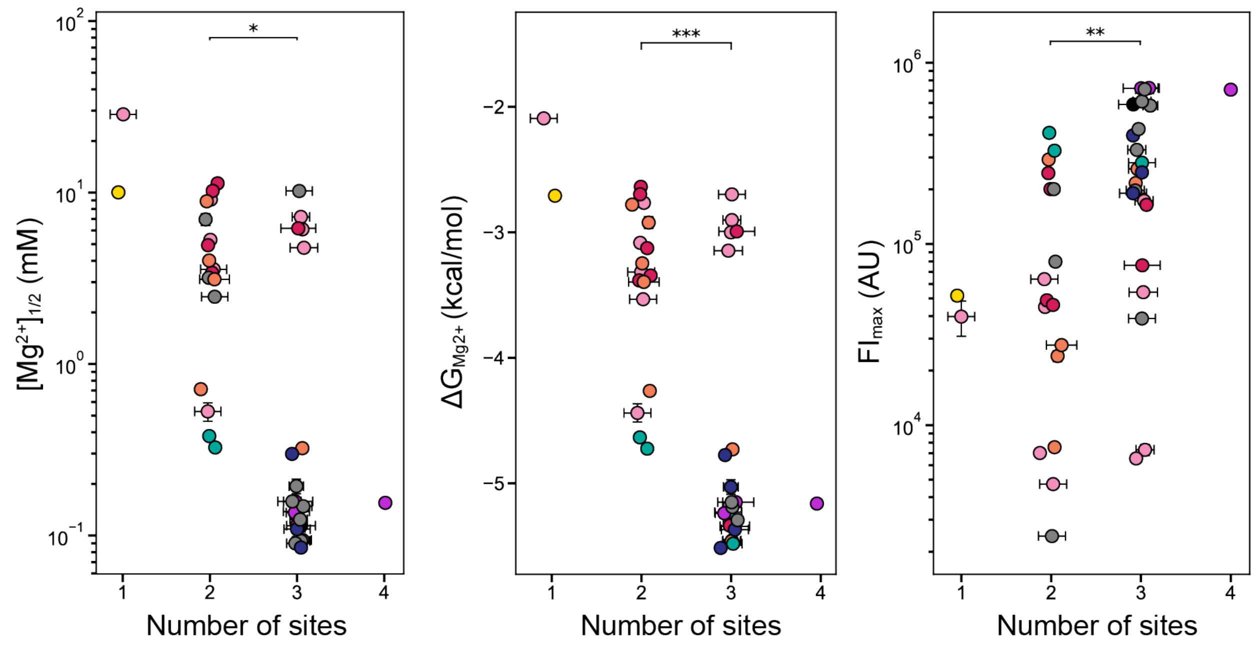


**Figure S11**. Metal binding site analysis. Jitter plots of (from left to right): fitted $[Mg^{2+}]_{1/2}$ values, $\Delta G_{Mg2+}$ (kcal/mole), and fitted $FI_{max}$ for constructs grouped by number of $Mg^{2+}$ sites.
